# Complexity and graded regulation of neuronal cell type-specific alternative splicing revealed by single-cell RNA sequencing

**DOI:** 10.1101/2021.01.27.428525

**Authors:** Huijuan Feng, Daniel F. Moakley, Shuonan Chen, Melissa G. McKenzie, Vilas Menon, Chaolin Zhang

## Abstract

The enormous neuronal cellular diversity in the mammalian brain, which is highly prototypical and organized in a hierarchical manner, is dictated by cell type-specific gene regulatory programs at the molecular level. Although prevalent in the brain, contribution of alternative splicing (AS) to the molecular diversity across neuronal cell types is just starting to emerge. Here we systematically investigated AS regulation across over 100 transcriptomically defined neuronal types of the adult mouse cortex using deep single cell RNA-sequencing (scRNA-seq) data. We found distinct splicing programs between glutamatergic and GABAergic neurons and between subclasses within each neuronal class, consisting of overlapping sets of alternative exons showing differential splicing at multiple hierarchical levels. Using an integrative approach, our analysis suggests that RNA-binding proteins (RBPs) Celf1/2, Mbnl2 and Khdrbs3 are preferentially expressed and more active in glutamatergic neurons, while Elavl2 and Qk are preferentially expressed and more active in GABAergic neurons. Importantly, these and additional RBPs also contribute to differential splicing between neuronal subclasses at multiple hierarchical levels, and some RBPs drive splicing dynamics that do not conform to the hierarchical structure defined by the transcriptional profiles. Thus, our results suggest graded regulation of AS across neuronal cell types, which provides a molecular mechanism orthogonal to, rather than downstream of, transcriptional regulation in specifying neuronal identity and function.

**Significance:** Alternative splicing (AS) is extensively used in the mammalian brain, but its contribution to the molecular and cellular diversity across neuronal cell types remains poorly understood. Through systematic and integrative analysis of AS regulation across over 100 transcriptomically defined cortical neuronal types, we found neuronal subclass-specific splicing regulatory programs consists of overlapping alternative exons showing differential splicing at multiple hierarchical levels. This graded AS regulation is controlled by unique combinations of RNA-binding proteins (RBPs). Importantly, these RBPs also drive splicing dynamics across neuronal cell types that do not conform to the hierarchical taxonomy established based on transcriptional profiles, suggesting that the graded AS regulation provides a molecular mechanism orthogonal to transcriptional regulation in specifying neuronal identity and function.

## Introduction

The structural and functional complexity of the mammalian cortex is determined by the enormous diversity of neuronal cell types and their connections that form intricate neural circuitries. Traditionally, neuronal cell types are defined by their morphological or electrophysiological properties, connectivity, or a small set of molecular markers (1–5). Recent advances in single-cell RNA sequencing (scRNA-seq) have enabled the unbiased discovery and characterization of neuronal cell types based on global transcriptional profiles (6–11). These studies demonstrated unprecedented power in revealing over 100 previously known and novel putative cell types. It has been suggested that these cell types can be organized with a hierarchical taxonomy consisting of glutamatergic (excitatory) neurons and GABAergic (inhibitory) interneurons as major neuronal classes. Among them, glutamatergic neurons are composed of five major branches, including L6 corticothalamic (CT), L5/6 near-projecting (NP), L5 pyramidal tract (PT), intratelencephalic (IT), and L6b subclasses, while GABAergic neurons can be divided based on their embryonic origins in the medial and caudal ganglionic eminences (MGE and CGE), which can be further divided into Pvalb and Sst subclasses for MGE neurons and Vip, Lamp5, Sncg, and Serpinf1 subclasses for CGE neurons (6). However, most of these transcriptional cell types remain poorly characterized.

One limitation in previous analysis of transcriptomic neuronal cell types is that those studies almost exclusively focused on the steady-state transcript level, while mammalian genes undergo multiple steps of post-transcriptional regulation which are essential for diversifying the final protein products in time and space. Alternative splicing (AS) of precursor mRNA (pre-mRNA) is a molecular mechanism to produce multiple transcript and protein isoforms through different combinations of exons (12). Transcriptomic analysis based on RNA-seq suggests that AS is ubiquitous, occurring in > 90% of multi-exon genes in human and other mammalian species (13, 14). The use of AS in the nervous system is particularly extensive, and previous studies have revealed distinct splicing profiles of brain tissues compared to non-neuronal organs (13–15), neurons compared to non-neuronal cells in the cortex (16, 17), central nervous system neurons compared to peripheral sensory neurons (18), as well as at different developmental stages in the cortex (18). While the functional significance of the majority of these neuron-specific alternative exons has yet to be demonstrated, there is no lack of examples in which individual alternative exons play critical roles in nervous system development and function, such as neuronal migration (19), axon outgrowth, pathfinding, and maturation (20–22), and synaptic formation and transmission (23, 24). In addition, emerging evidence suggests global splicing differences between major excitatory and inhibitory neuronal cell types from recent bulk RNA-seq analysis of specific cell types or ribosome-associated transcriptomes purified from transgenic mice (25, 26). Therefore, elucidating cell type-specific splicing regulation is a key step toward understanding the underpinnings of neuronal cell type diversity and function.

Cell type-specific AS is largely controlled by RNA-binding proteins (RBPs) that recognize specific regulatory sequences embedded in pre-mRNA transcripts. Technological advances have also made it possible to define the comprehensive RBP target networks by integrating global splicing profiles upon RBP depletion and genome-wide maps of protein-RNA interactions, as demonstrated by multiple studies, including our own (27, 28). For instance, RBPs specifically expressed or enriched in neurons, such as Nova, Rbfox, Ptbp2, nElavl, Srrm4 (nSR100) and Mbnl2, have been demonstrated to regulate AS of numerous neuronal transcripts [reviewed in refs. (29–31)]. In addition, KH domain-containing proteins Khdrbs2 (Slm1) and Khdrbs3 (Slm2) were recently shown to be preferentially expressed in pyramidal cells over Pvalb interneurons in hippocampus to regulate splicing of a highly specific set of transcripts involved in excitatory synaptic formation (32, 33).

Despite this exciting progress, the extent to which AS contributes to the diversity of the wide range of neuronal cell types and the underlying regulatory mechanisms has not been systematically investigated. In the present study, we took advantage of deep scRNA-seq data of adult mouse neocortex with whole transcript coverage to identify differentially spliced (DS) alternative exons in major neuronal classes and subclasses at different hierarchical levels. By integrating *de novo* motif analysis, RBP expression profiles, their target networks and position-dependent RNA-maps, our analysis revealed several major regulators of the neuronal cell type-specific AS and “graded”, combinatorial regulatory programs that drive the complex landscape of splicing dynamics across cell types.

## Results

### Major neuronal cell types revealed by single-cell splicing profiles

To study neuronal cell type-specific AS on a global scale, we utilized the two sets of scRNA-seq data derived from adult mouse cortex with deep and full length transcript coverage, generated by the Allen Institute for Brain Science (6, 7) (denoted as the Tasic 2016 and Tasic 2018 datasets hereafter). The Tasic 2016 dataset includes 1,809 cells from the primary visual cortex (VISp), while the Tasic 2018 dataset is composed of 15,413 cells from VISp and 10,068 cells from anterior lateral motor cortex (ALM). In our analysis, we used the 1,424 and 21,154 core cells that were unambiguously assigned to different transcriptional cell types as defined in the two original studies (totaling 14 billion reads and 54 billion read pairs, respectively), and quantified the inclusion level (percent spliced in or PSI, denoted Ψ) of over 16,000 cassette exons using the Quantas pipeline (16). Overall, a comparable number of exons have sufficient exon junction reads (i.e., ≥20) for splicing quantification in glutamatergic and GABAergic neuronal populations in each dataset, while a lower number of exons were quantifiable in non-neuronal cells (median=2,328, 2,114, and 813 for glutamatergic neurons, GABAergic neurons and non-neuronal cells, respectively, in Tasic 2016 dataset and 2,176, 2,218, and 1,128 in Tasic 2018 dataset; Fig. S1 and Table S1), which is consistent with lower RNA yield in non-neuronal cells. Given its much larger number of cells, we primarily focused on the Tasic 2018 dataset unless explicitly specified, and integrated results from Tasic 2016 dataset when appropriate.

We first asked whether single-cell splicing profiles can be used to infer neuronal cell types in the cortex. To this end, we performed t-distributed neighbor embedding (t-SNE) analysis and clustered cells using single-cell splicing profiles (Materials and Methods). For the Tasic 2018 dataset, we identified nine major clusters (Fig. 1A). When compared to the 133 transcriptional cell types defined originally based on gene expression profiles, which consist of 56 glutamatergic, 61 GABAergic and 16 non-neuronal cell types, the clusters defined by splicing profiles clearly separated glutamatergic (Clusters 1-4), GABAergic (Clusters 5-7) and non-neuronal cells (clusters 8-9). Different subclasses within each of these major cell classes can also be distinguished, although they overlap in the t-SNE plots. GABAergic neurons form the three major clusters, corresponding to Vip and Lamp5-Sncg-Serpinf1 (Cluster 5), Sst (Cluster 6) and Pvalb (Cluster 7) subclasses, respectively. In the glutamatergic neuronal population, cluster 1 clearly represents VISp-specific IT neurons from layers 2/3 and layer 6 (Fig. 1B), while the other subclasses cannot be distinguished at this resolution. The non-neuronal cells are clearly separated into two major clusters with one corresponding to astrocytes and oligodendrocytes (cluster 8) and the other representing endothelial, vascular and microglia cells which do not arise from the neuroectoderm lineage (Cluster 9). Similar observations were made from the Tasic 2016 dataset (Fig. S2A-C and Table S1). When we colored the single cells by cortical regions, we observed several dense areas with ALM/VISp-derived cells in the glutamatergic neuron clusters but no obvious regional aggregation of cells in the GABAergic neuron clusters (Fig. 1A), which is similar to the observation by Tasic et al. based on gene expression profiles (6). These data suggest that the global splicing profile at the single-cell level can clearly define the two primary neuronal classes, and to some extent, several major subclasses in the cortex.

**Figure 1:**
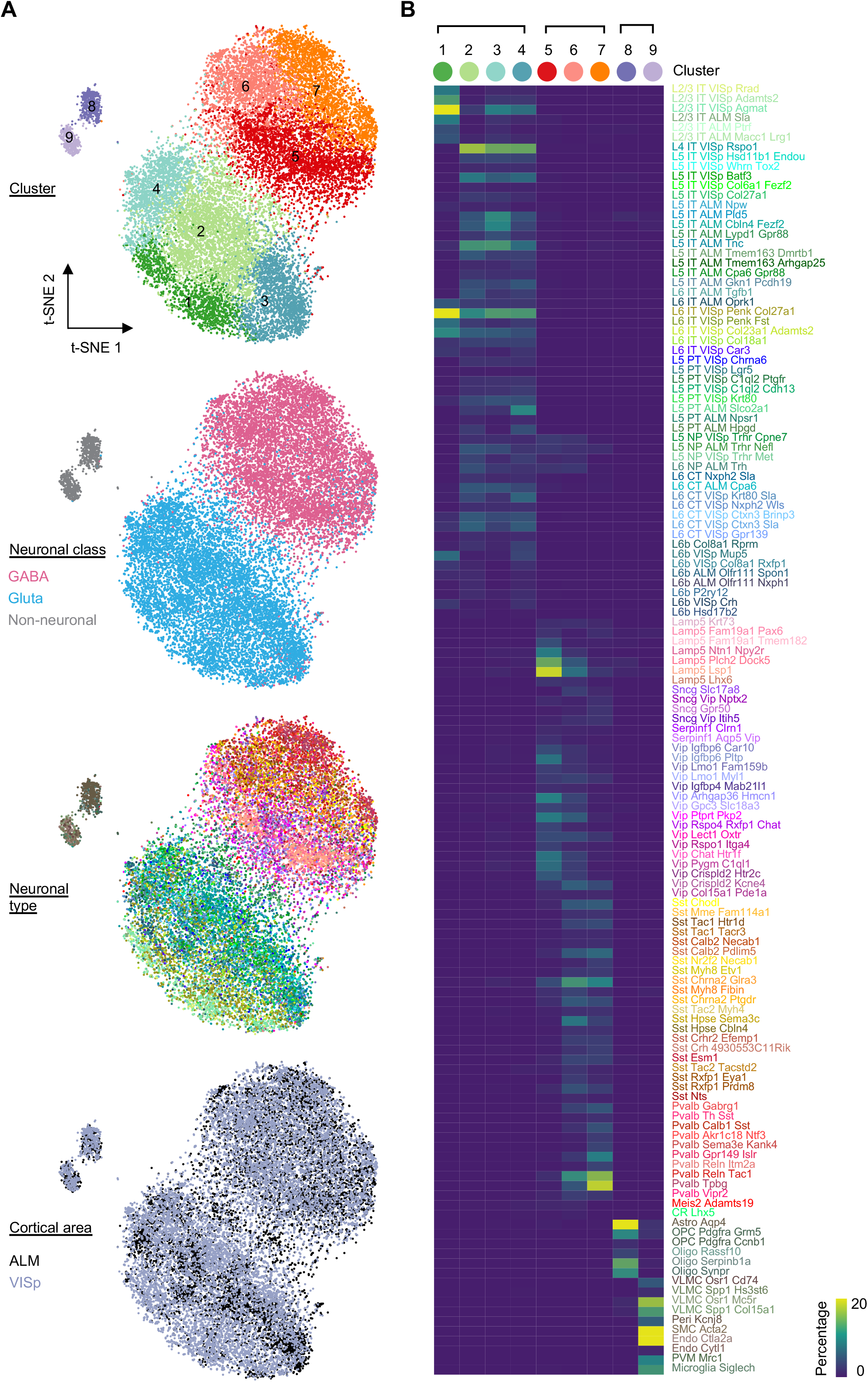
Clustering analysis of single cell splicing profiles reveals major neuronal and glial cell classes. **A**, Two-dimensional t-distributed stochastic neighbor embedding (t-SNE) plots of 19,853 core cells from the Tasic 2018 scRNA-seq dataset. Cells are colored by cluster based on the splicing profile, cortical area of the cell, neuronal class, and neuronal types defined based on gene expression profile from the original paper. **B**, Heatmap showing the overlap between the clusters defined based on splicing profiles and the original neuronal cell types defined based on expression profiles.

### Differential splicing between glutamatergic and GABAergic neurons

Having demonstrated the neuronal cell type specificity carried in the splicing profile, we next sought to identify alternative exons differentially regulated between different neuronal classes and subclasses. To optimize our analysis pipeline, we initially focused on the two primary neuronal classes, glutamatergic and GABAergic neurons and detected 937 and 901 DS cassette exons in the Tasic 2016 and Tasic 2018 datasets, respectively (changes in PSI or |ΔΨ|≥0.1, false discovery rate (FDR)≤0.05; cells from ALM and VISp were aggregated together for this analysis; Materials and Methods). With more stringent criteria (ΔΨ|≥0.2), 342 and 269 cassette exons show differential splicing, totaling 469 DS exons from 368 genes used for additional characterization below (Table S2). DS exons detected in the two datasets largely overlap with 142 exons in common, all of which show splicing differences in the same direction (Fig. 2A). Quantitatively, the ΔΨ values of the 469 unique DS exons are correlated well between the two datasets (Pearson’s correlation R=0.44, Fig. 2B and Table S2).

**Figure 2:**
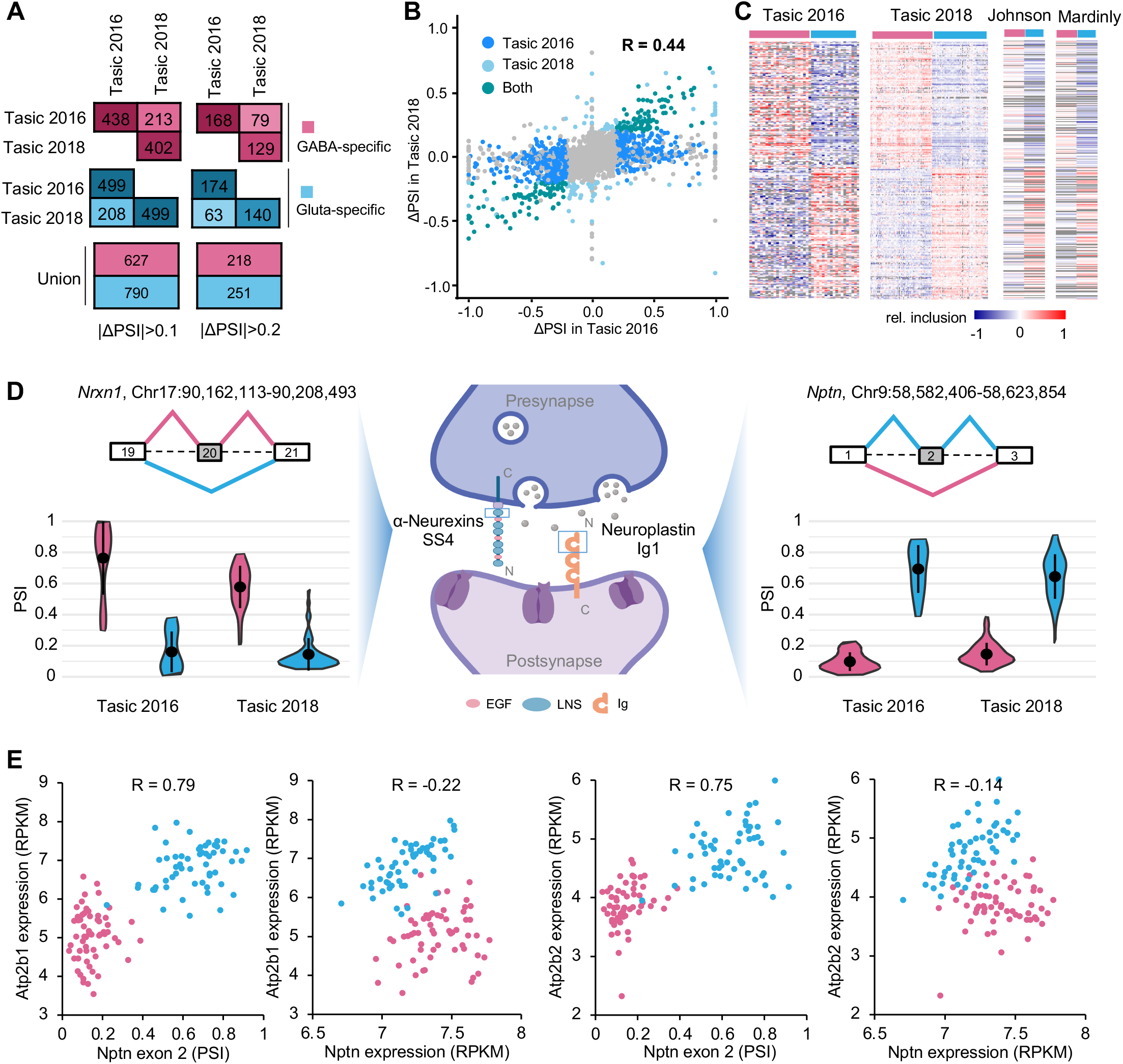
A robust set of DS cassette exons between glutamatergic and GABAergic neurons identified from scRNA-seq data. **A**, Number of DS exons with higher inclusion in either GABAergic neurons (GABAergic-specific exons, pink) or glutamatergic neurons (glutamatergic-specific exons, blue) detected in Tasic 2016 and Tasic 2018 datasets. **B**, Correlation of ΔΨ between GABAergic and glutamatergic neurons for all cassette exons estimated from Tasic 2016 and Tasic 2018 datasets. Dark blue points represent the DS exons (ΔΨ≥0.2) only detected in Tasic 2016, light blue points represent the DS exons only detected in Tasic 2018, green points represent the overlap of DS exons detected in both datasets and grey points represent the other non-DS exons. The Pearson correlation is indicated. **C**, Heatmaps showing exon inclusion level (with median subtraction) of DS exons in the two scRNA-seq datasets and two independent bulk RNA-seq datasets. Only DS exons with quantifiable exon inclusion in ≥50% of neuronal cell types in both Tasic 2016 and 2018 datasets were included for visualization. **D**, Examples of DS exons in synaptic-related genes *Nrxn1* and *Nptn* showing GABAergic or glutamatergic-specific inclusion, respectively. The violin plots show the distribution of PSI in single cells of GABAergic and glutamatergic neuronal classes. Median and the interquantile range are indicated. **E,** *Nptn* exon 2 inclusion but not gene expression is correlated with Pmca (encoded by Atp2b1 and Atp2b2) expression. Glutamatergic and GABAergic neuronal cell types are colored in blue and pink, respectively. Pearson’s correlation is indicated in each scatter plot.

To assess potential regional differences, we also analyzed cells from ALM and VISp in the Tasic 2018 dataset separately. Splicing differences between glutamatergic and GABAergic neurons detected in the two regions are consistent in directions and also well correlated quantitatively (R=0.38, Fig. S3A,B and Table S2). Therefore, the differential splicing regulatory program between glutamatergic and GABAergic neurons appears to be largely shared across the two cortical regions, justifying the aggregation of cells across regions in our analysis.

To further validate the reliability of DS exons detected in scRNA-seq data, we analyzed three bulk RNA-seq datasets of purified glutamatergic and GABAergic neurons (34–36) (Table S1). A total of 320 DS exons were detected in one or more bulk RNA-seq dataset, with an average of 147 exons detected in individual datasets (Table S3). These exons have large overlap with the 469 DS exons we detected in scRNA-seq data with 160 exons in common and splicing differences in the same direction (Fig. 2C).

Compared to all cassette exons, the DS exons show characteristics of highly regulated, potentially functional alternative exons, with a higher percentage of in-frame coding exons (>75% vs. 54%) and more conserved AS patterns in human and/or rat (>49% vs. 24%). In addition, a larger proportion of DS exons show neuron-specific splicing when comparing neuronal vs non-neuronal cells in the cortex (>24% vs. 6%) and dynamic splicing switches at different developmental stages (~60% vs. 12%) (Fig. S4A; Tables S2 and S3). Gene ontology (GO) analysis using the 368 genes containing DS exons revealed significant enrichment of genes under three major GO categories including synapse, cell projection and ion channel activity (Fig. S4B,C), in agreement with previous observations (25, 26). Therefore, our computational pipeline for scRNA-seq data is able to reliabily detect differential splicing between distinct neuronal populations and a majority of these exons are missed by bulk RNA-seq data, suggesting the superior sensitivity of our analysis when applied to scRNA-seq data sets.

As individual examples, our analysis detected GABAergic neuron-specific inclusion of the well-studied *Nrxn1* exon 20 (also known as alternatively spliced site 4, or SS4), which encodes a peptide overlapping with the sixth LNS domain of the Neurexin 1 protein (Fig. 2D, left and middle panels). AS of this exon is known to be neuronal activity-dependent and may contribute to neuron-type specific synaptic development by modulating its postsynaptic binding partners (23, 37). As a second, less characterized example, we identified *Nptn* exon 2 to be highly included in glutamatergic neurons but predominantly skipped in GABAergic neurons (Fig. 2D middle and right panels). *Nptn* encodes cell adhesion molecule that mediates formation and stabilization of excitatory synapses, activity-dependent long-term synaptic plasticity, as well as excitatory/inhibitory balance (38). Specific depletion of Nptn in glutamatergic neurons in mice results in altered neuronal activity and behavioral deficits by reducing the level of plasma membrane calcium ATPase (Pmca) genes Atp2b1 and Atp2b2 and thus increasing intracellular Ca^2+^ concentration (39). Inclusion of exon 2 generates a longer isoform Np65 containing three Ig domains, which was demonstrated to be brain- and neuron-specific and localized between pre- and postsynaptic components of excitatory synapses (38). Interestingly, our analysis suggests that the inclusion of exon 2, but not the overall *Nptn* mRNA level, is strongly correlated with Pmca gene expression across neuronal cell types (R= 0.79 vs. −0.22 for *Atp2b1*; R=0.75 vs. −0.14 for *Atp2b2*; Fig. 2E). Therefore, Np65 might be the major contributor that regulates distinct Pmca expression between glutamatergic and GABAergic neurons.

### Neuronal subclass-specific splicing programs at different hierarchical levels

Compared to bulk RNA-seq data derived from a priori defined neuronal cell types, scRNA-seq data with in-depth sampling of cells such as the Tasic 2018 dataset offers the unique opportunity to interrogate differential splicing between neuronal subclasses defined at different hierarchical levels. Our t-SNE analysis of splicing profiles showed aggregation of single cells reflecting subclasses of glutamatergic and GABAergic neuronal populations, although they overlap in the low-dimensional representation afforded by t-SNE (Fig. 1A). This overlap could be partly due to insufficient read coverage that resulted in the uncertainty in the single cell splicing profiles. To reduce the uncertainty of splicing quantification, we performed an unbiased hierarchical clustering analysis of the splicing profiles of 55 glutamatergic and 60 GABAergic transcriptional cell types, respectively, by pooling the single cells assigned to each neuronal cell type (Fig. 3A). This analysis revealed clear separation of CGE- and MGE-originating GABAergic neurons, as well as Vip, Sncg, Lamp5, and Serpinf1 subclasses in the CGE cluster and Pvalb and Sst subclasses in the MGE cluster. The glutamatergic neurons are mainly divided by projection subtypes with IT neurons constituting the largest branch, followed by PT, NP and L6b neuronal subclasses, while CT neurons are distributed across the branches (Fig. 3B). Within the IT branch, we observed higher similarity between L2/3 IT and L6 IT neurons as compared to L4/5 neurons (Fig. 3B). These observations confirmed that the molecular diversity at the splicing level not only contributes to the distinction of glutamatergic vs. GABAergic neuronal classes, but also the delineation of subclasses in each population at multiple hierarchical levels.

**Figure 3:**
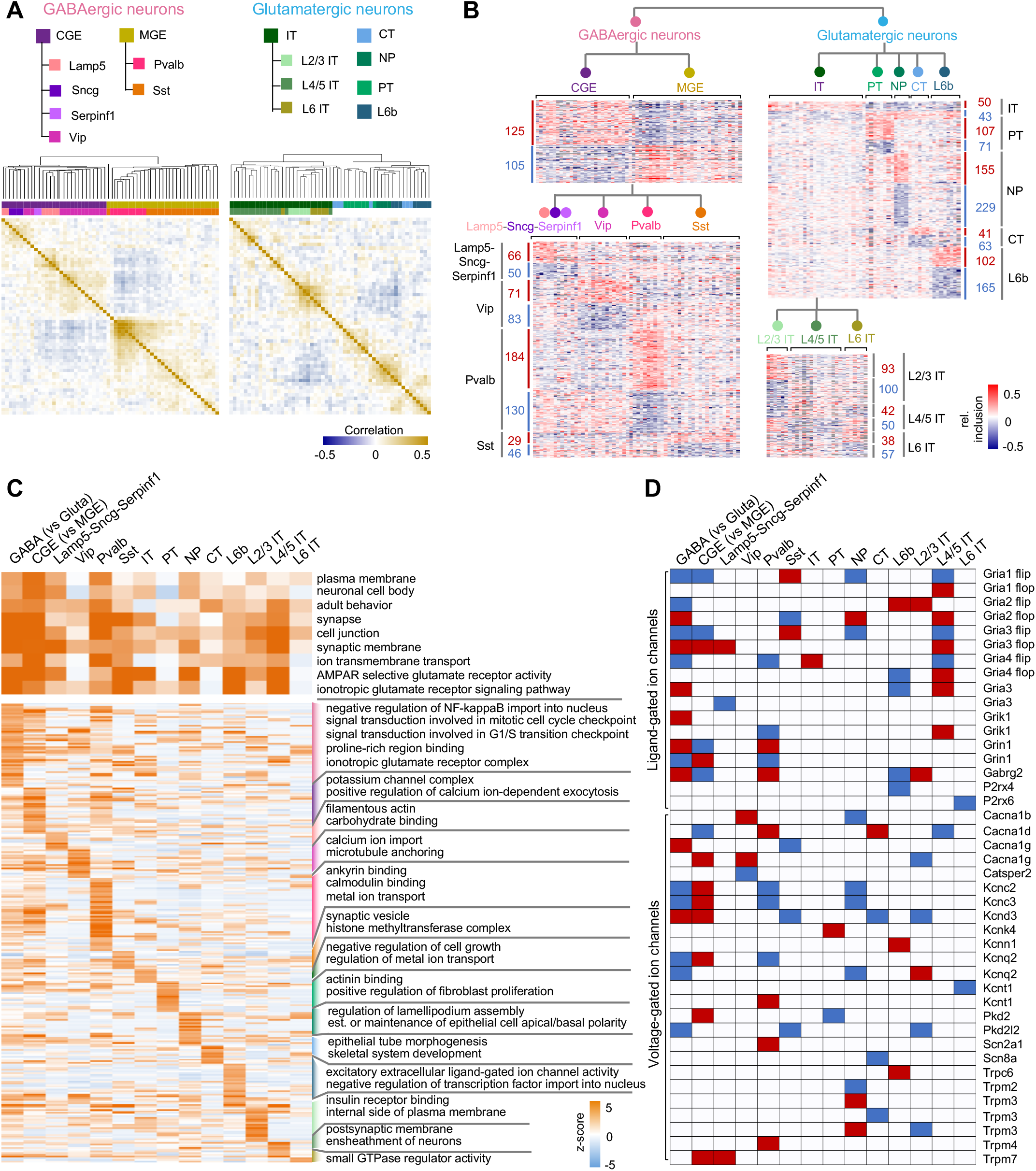
Neuronal subclass-specific splicing identified at different hierarchical levels. **A**, Heatmap showing the pairwise Pearson's correlation coefficient of splicing profiles across neuronal cell types within the GABAergic (left) or glutamatergic (right) neuronal class. In each panel, exons quantifiable in ≥50% of cell types in the respective neuronal class with standard deviation (SD) ≥0.1 were used for analysis. Cell types are ordered by hierarchical clustering with the dendrogram tree shown at the top. Cell types are colored by their developmental origin (MGE or CGE) and major subclasses (Pvalb, Sst, Vip, Lamp5, Sncg, and Serpinf1 for GABAergic neurons and IT, CT, PT, NP and L6b for glutamatergic neurons; IT subclasses further divided by layer. **B**, Heatmaps showing relative exon inclusion level (with median subtraction) of DS exons between neuronal subclasses detected at different hierarchical levels. Each row represents an exon and each column represents a cell type. The number of DS exons in each comparison is labeled. **C,** Gene ontology (GO) terms associdated with DS exons contrasting neuronal classes or subclasses at different hierarchical levels. Shared GO terms by multiple comparisons (i.e., significant in ≥5 comparisons) are shown at the top, while GO terms enriched in specific subclasses are shown at the bottom. GO term enrichment z-scores are shown in the heatmap with color scale indicated.

Consistent with the unsupervised clustering analysis, we identified robust sets of DS exons by contrasting neuronal subclasses defined at different hierarchical levels and these exons also showed characteristics of highly regulated AS exons (Fig. S5). For the GABAergic neuronal types, 230 exons are differentially spliced between MGE and CGE-originating neurons, and between 75 and 314 exons show differential splicing when the four major subclasses Pvalb, Sst, Vip, and Lamp5-Sncg-Serpinf1 neurons were compared (ΔΨ|≥0.1, FDR≤0.05; Fig. 3B and Table S4). Similarly, between 93 and 384 DS exons were identified between major glutamatergic subclasses including IT, PT, NP, CT and L6b neurons, with the largest number observed from NP neurons (Fig. 3B and Table S4). In addition, IT neurons represent a heterogenous group, and subsets of exons show distinct splicing patterns depending on the cortical layers from which they are derived.

To inform the potential function associated with neuron class or subclass-specific AS at different hierarchical levels, we examined GO terms enriched in lists of genes containing DS exons for each neuron subclass. We found that terms related to cell-cell adhesion, synapse, and ion channel/receptor activity are shared by multiple subclasses, suggesting that an enormous AS-mediated diversity is required for these genes related to fundamental neuronal properties (Fig. 3C and Table S5). In particular, we found that both ligand-gated and voltage-gated ion channel genes frequently contain specific isoforms enriched or depleted in multiple subclasses (Fig. 3D). On the other hand, our analysis also identified GO terms that are specifically enriched in certain subclasses, such as “excitatory extracellular ligand-gated ion channel activity” in L6b neurons and “calmodulin binding” and “metal ion transport” in Pvalb neurons (Fig. 3C and Table S5).

In accordance with the complex splicing regulation of ion channel gene families, comparison between lists of DS exons contrasting neuronal classes and subclasses at different hierarchical levels showed interesting overlaps (Fig. 4A). In particular, we noticed a significant overlap between exons differentially spliced in GABAergic vs. glutamatergic neurons and those between NP vs. other subclasses of glutamatergic neurons (p=1e-151; Fisher’s exact test), with similar splicing patterns in GABAergic and NP neurons. One major characteristic feature shared between GABAergic interneurons and NP neurons, but distinct from the other types of glutamatergic neurons, is the range of projection. To investigate whether a shared splicing program underlies this property, we took advantage of a subclass of L6b neurons which also have local projections (6) and identified DS exons between local vs. long projecting L6b neurons. Comparison of the three lists of DS exons allowed us to identify 41 common exons with essentially all exons showing consistent splicing patterns with respect to projection types (Fig. 4B,C and Table S6). Genes containing these exons have functions associated with dendrite, synapse and cell junctions, including multiple genes such as *Tsc2*, *Arhgap44* and *Macf1*, which were previously demonstrated to play a role in axon outgrowth and pathfinding (40–42). In addition, nine genes were shown to undergo dynamic AS switches during axonogenesis (21) (odds ratio=11, p=8.2e-7, Fisher’s exact test) (Fig. 4D). While further experimentation is required, these data suggest a splicing program correlated with and potentially contributing to near vs. long range projection. Taken together, the hierarchical analysis suggests a complex landscape of AS contributing to the molecular diversity across neuronal cell types.

**Figure 4:**
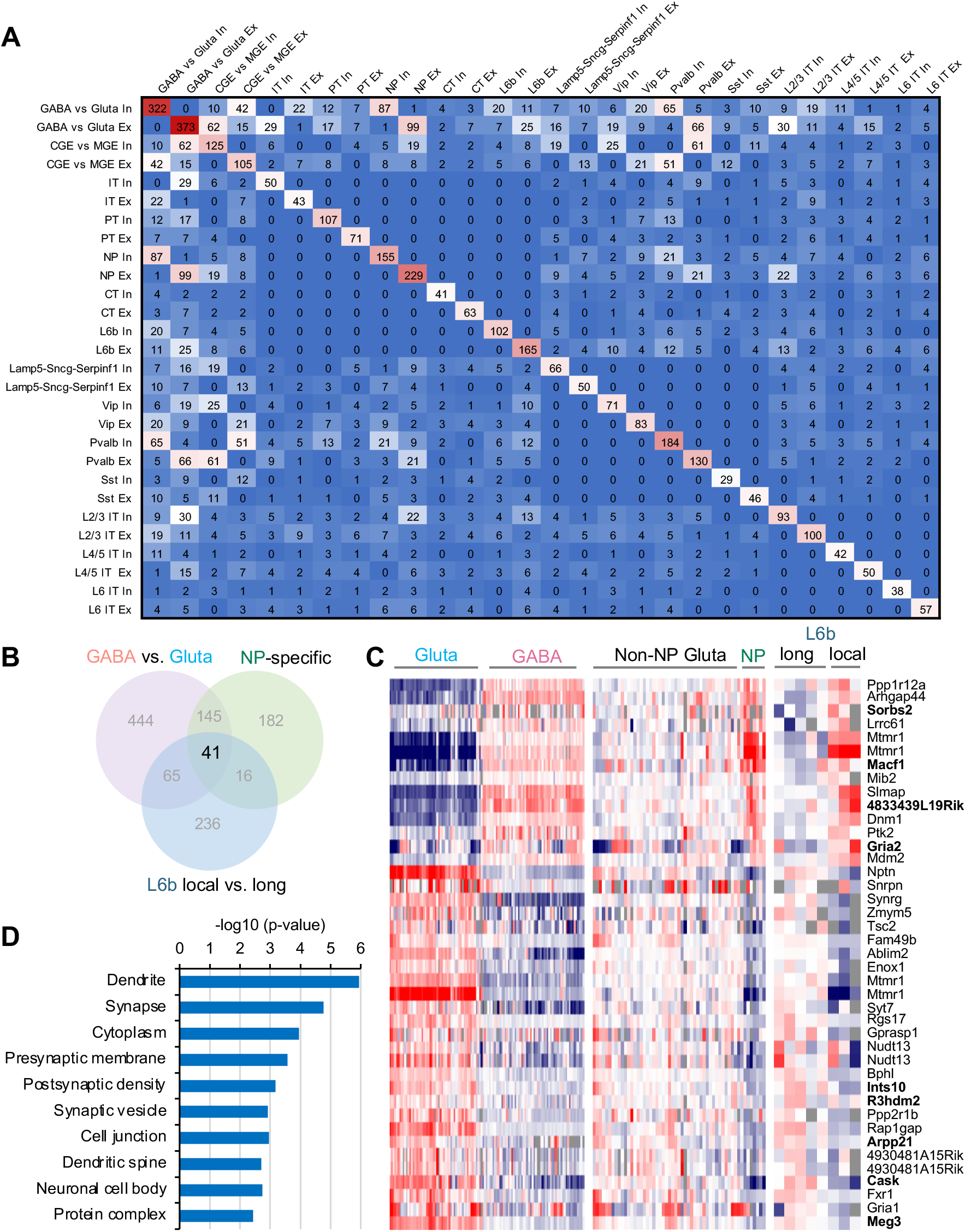
Graded regulation of splicing across neuronal cell types. **A**, Pairwise comparison showing overlap of DS exons detected at different hierarchical levels. Exons with neuronal class or subclass-specific inclusion and skipping are shown separately. **B**, Overlap of DS exons identified in comparison of neuronal subclasses at different hierarchical levels potentially associated with range of neuronal projection. **C,D**, Splicing profile (C) and associated GO terms (D) for 41 commons DS exons identified in (B). Exons in genes showing axonogenesis-dependent AS switches(21) are highlighted in bold in (C).

### RBPs regulating differential splicing between glutamatergic and GABAergic neurons

We next asked how the distinct splicing programs across neuronal cell types are controlled. Unbiased clustering analysis of RBP expression clearly segregated glutamatergic and GABAergic neurons, implying their potential role in regulating differential splicing between neuronal classes (Fig. S6 and Table S7). Differential expression analysis identified 17 RBPs (e.g., Celf2/5, Khdrbs2/3, Rbfox3, and Mbnl2) showing higher expression in glutamatergic neurons and 24 RBPs (e.g., Qk, Elavl2, Rbmx, and Upf3b) showing higher expression in GABAergic neurons (FDR≤0.05, fold change≥1.5; Fig. S7 and Table S7). In addition, we identified 78 RBPs that are differentially expressed in specific neuronal subclasses within glutamatergic or GABAergic neuronal classes at different hierarchical levels (Fig. S7 and Table S7).

To identify regulators of differential splicing between glutamatergic and GABAergic neurons, we began with an unbiased *de novo* motif analysis by searching for k-mers (k=4, 5, 6) enriched in the glutamatergic or GABAergic neuron-specific alternative exons and flanking intronic sequences (within 200 nucleotide region). This analysis identified 78 4-mers, 71 5-mers and 56 6-mers that are enriched in or around glutamatergic or GABAergic neuron-specific exons (FDR≤0.05, hypergeometric test; Table S8). We clustered the significant k-mers to generate a k-mer graph for visualization based on their sequence similarity as well as position-dependent enrichment patterns since many RBPs activate or repress exon inclusion depending on where they bind (frequently activate exon inclusion when binding to the downstream and repress exon inclusion when binding to the alternative exon or upstream; Materials and Methods). The enriched k-mers can be organized into several clusters resembling RBP binding sites known to regulate AS in neurons (Fig. 5). The largest cluster consists of UG or GU-rich sequences that are recognized by the Celf family RBPs; these k-mers are enriched in the downstream intron for exons preferentially included in glutamatergic neurons and in the upstream intron for exons preferentially included in GABAergic neurons. The position-dependent enrichment pattern is most consistent with the higher expression of Celf2 in glutamatergic neurons (and to some extent, also Celf1, which has significant albeit more moderate differences), so that Celf1/2-dependent splicing activation by binding to the downstream intron results in preferential exon inclusion in glutamatergic neurons while Celf1/2-dependent splicing repression by binding to the upstream intron results in preferential exon skipping in glutamatergic neurons (i.e., preferential exon inclusion in GABAergic neurons). Similar is true for k-mers containing YGCY tetramers (Y=C/U), representing the consensus binding motif of Mbnl proteins, especially Mbnl2, which is also more highly expressed in glutamatergic neurons. In contrast, a cluster of U-or AU-rich k-mers shows enrichment in the downstream intron of GABAergic neuron-specific exons and the upstream intron of glutamatergic-specific exons. This enrichment pattern is most consistent with Elavl2, which is more highly expressed in GABAergic neurons, so that its binding in the downstream intron results in splicing activation and preferential exon inclusion in GABAergic neurons, while its binding in the upstream intron results in splicing repression and preferential exon skipping in GABAergic neurons (i.e., preferential exon inclusion in glutamatergic neurons). The other two clusters contain UWAA (W=U/A) and C-rich motifs, which are known as known as the binding consensus of Khdrbs and Pcbp families, respectively.

**Figure 5:**
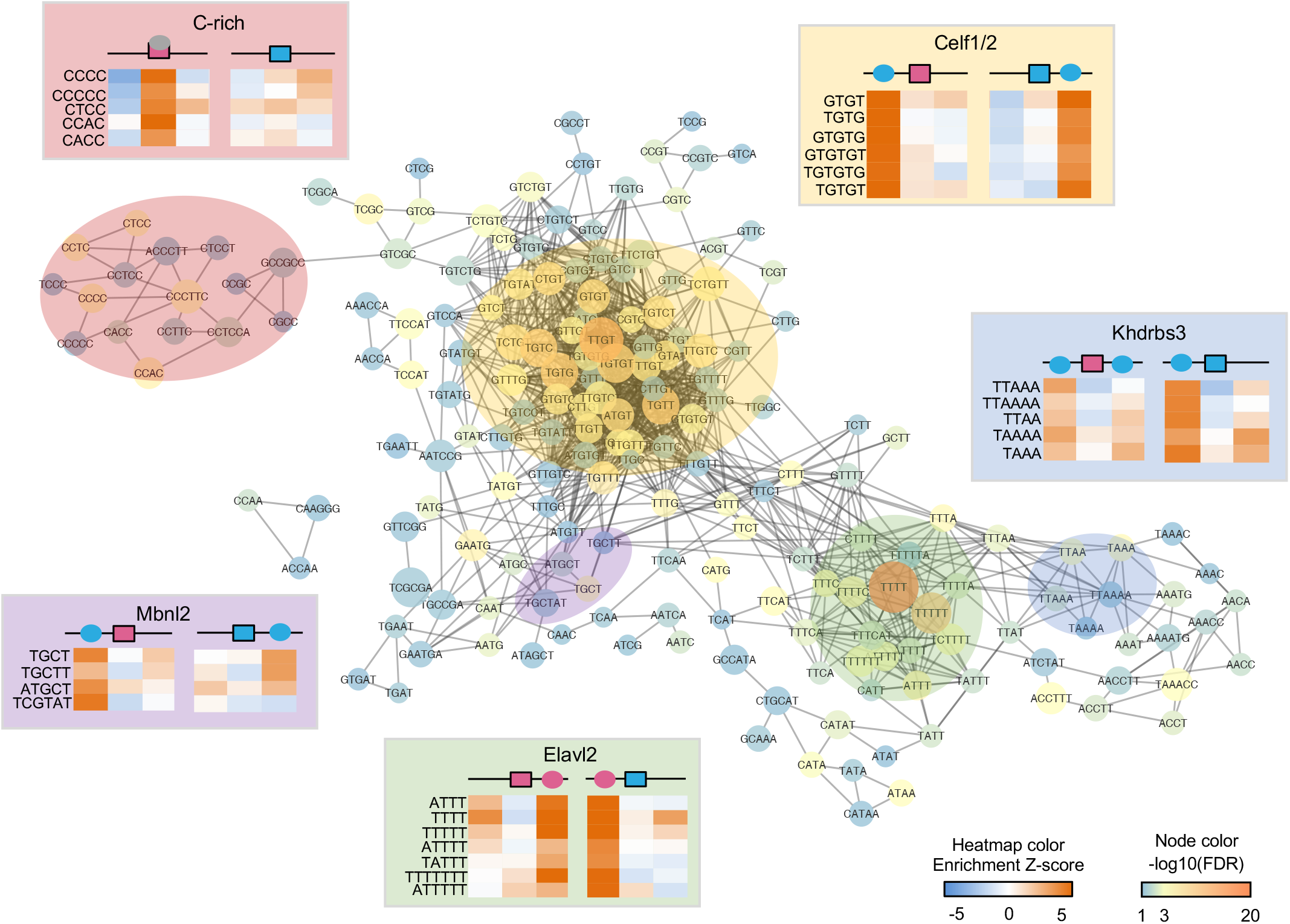
*De novo* motif analysis of DS exons with glutamatergic or GABAergic neuron-specific splicing. The graph shows significantly enriched k-mers (k= 4-6) in either glutamatergic or GABAergic neuron-specific exons or their flanking intronic sequences. K-mers with similar sequences and position-dependent enrichment patterns are connected. Nodes are colored by the FDR of the most significant region. Five clusters of k-mers (shaded circles) are highlighted. For each cluster, the most likely cognate RBP (when available) is indicated and the position-dependent enrichment patterns of representative k-mers are also shown using a heatmap. Blue and pink colors represent higher RBP expression or exon inclusion in glutamatergic and GABAergic neurons, respectively.

To further establish that differential RBP expression directly contributes to the distinct splicing patterns between glutamatergic and GABAergic neurons, we examined splicing targets of several neuronal RBP families as determined by RBP depletion followed by RNA-seq or, whenever possible, integrative modeling of multiple types of data including RBP-dependent splicing and direct protein-RNA interactions, as we described previously (18, 27) (Table S9). Indeed, for RBPs with higher expression in GABAergic neurons (Elavl2 and Qk), exons with RBP-dependent inclusion are preferentially included in GABAergic neurons, while exons with RBP-dependent repression are preferentially included in glutamatergic neurons, consistent with the notion that higher activity of these RBPs in GABAergic neurons drives a GABAergic neuron-specific splicing profile. The opposite is true for Mbnl2 and Celf with higher expression in glutamatergic neurons. Khdrbs3 is known to mostly repress exon inclusion, consistent with the preferential skipping we see of its target exons in glutamatergic neurons (Fig. 6A). In contrast, Rbfox, Ptbp and Nova do not seem to have a systematic impact on the differential splicing between glutamatergic and GABAergic neurons, although Rbfox3 is more highly expressed in glutamatergic neurons. When we projected splicing profiles into the low dimensional space using principal component analysis (PCA), we observed shifts of splicing profiles towards GABAergic neurons upon depletion of Mbnl1/2, Celf1 and Khdrbs3, and shifts towards glutamatergic neurons upon depletion of Qk (no RNA-seq data with neuronal Elavl depletion), suggesting the global impact of these RBPs (Fig. 6B). Our analyses suggest that a combination of RBPs preferentially expressed in glutamatergic (Celf1/2, Mbnl2 and Khdrbs3) and GABAergic neurons (Elavl2 and Qk) together control the distinct splicing programs in the two neuronal classes.

**Figure 6:**
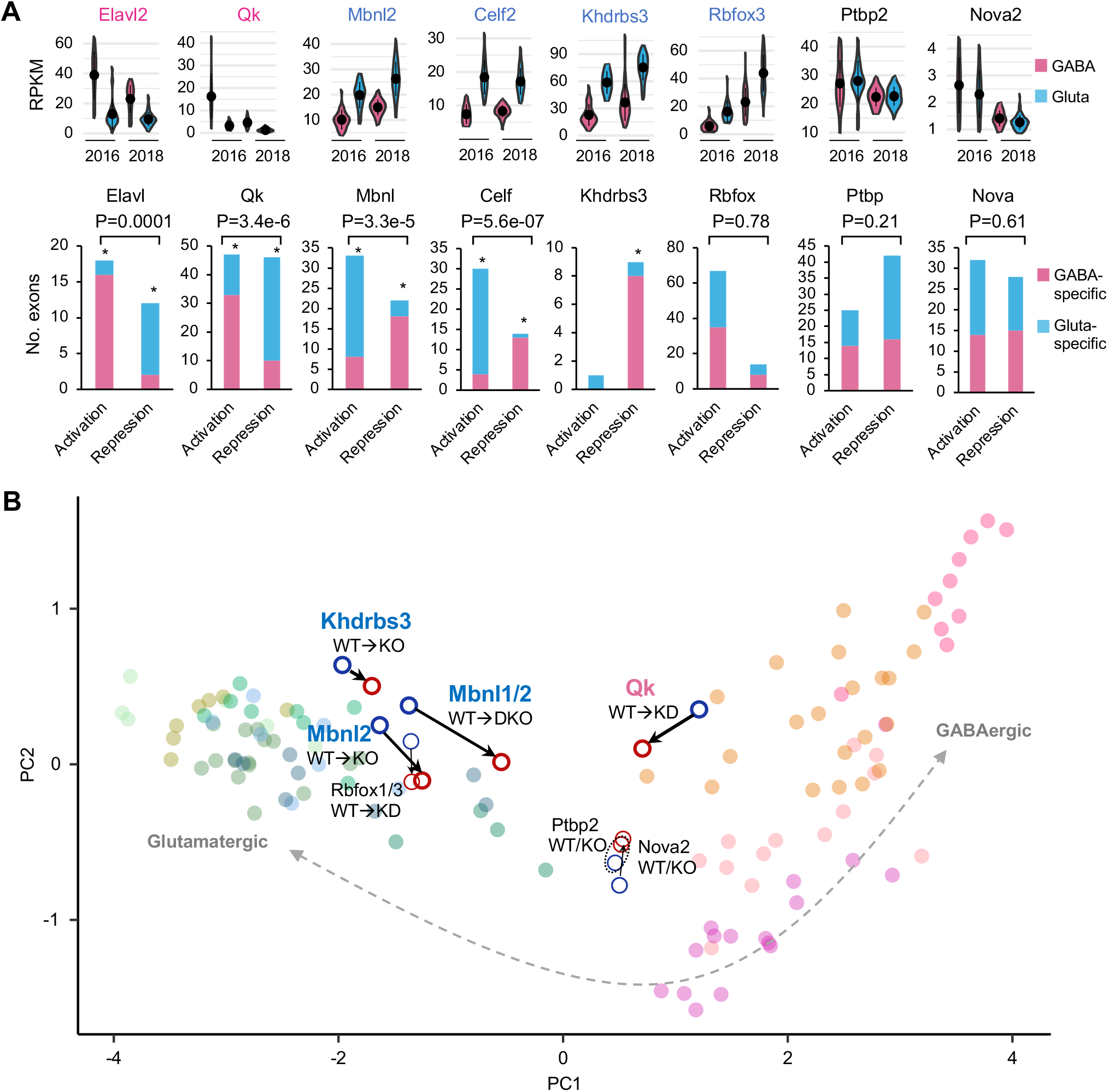
RBPs regulating differential splicing between glutamatergic and GABAergic neurons. **A**, Exons regulated by several RBP or RBP families showing glutamatergic or GABAergic neuron-specific exon inclusion. For each family, the representative RBP with differential expression between glutamatergic and GABAergic neurons estimated from Tasic 2016 and 2018 datasets is shown at the top. Note for Elavl2 and Qk with preferential expression in GABAergic neurons, exon activation or repression leads to a GABAergic neuron-like splicing profile, while for Mbnl2, Celf2 and Khdrbs3 with preferential expression in glutamatergic neurons, exon activation or repression leads to a glutamatergic neuron-like splicing profile. An asterisk indicates significant bias towards glutamatergic or GABAergic neuron-specific inclusion for exons activated or repressed by the respective RBP, as compared to all DS exons between glutamatergic and GABAergic neurons (p<0.05). Exons activated by an RBP and those repressed by the RBP are also compared and the resulting p-value is shown at the top. **B**, A global impact of RBP depletion on the shift of splicing profile towards glutamatergic or GABAergic neurons. In the scatter plot, each dot is a sample, which is projected into the two-dimensional space defined by the first two principal components of splicing profiles of glutamatergic or GABAergic neuronal cell types.

It is particularly intriguing that that about 20% of Mbnl-regulated exons show differential splicing between glutamatergic and GABAergic neurons (12% of DS exons of the two neuronal classes), presumably due to the higher expression and activity of Mbnl2 in glutamatergic neurons. The importance of Mbnl2 in regulating differential splicing between glutamatergic and GABAergic neurons has not been characterized, but is further supported by several lines of evidence in our analysis. The direct regulation of the DS exons by Mbnl2 is not only supported by position-dependent binding sites evident from motif analysis, but also from Mbnl2 binding footprints mapped by CLIP (Fig. 7A). In addition, 44% of DS exons between glutamatergic and GABAergic neurons show developmental splicing switches observed in mouse cortex, and exons with late splicing switches between P4 and P15 are particularly enriched, consistent with Mbnl2 as a major regulator that promotes the mature splicing program in the cortex (18, 43) (Fig. 7B). Strikingly, among DS exons regulated by Mbnl2, the differential splicing pattern between glutamatergic and GABAergic neurons is very similar to that between adult and embryonic cortices, and can be explained by Mbnl2-dependent activation or repression (Fig. 7C). As a specific example, exon 8 of metabotropic glutamate receptor 5 gene (*Grm5*, also known as mGluR5), which encodes 32 amino acids in the C-terminal region, is more highly included in glutamatergic over GABAergic neurons (PSI=0.81 vs. 0.41; Fig. 7D,E). Inclusion of this exon is strongly correlated with Mbnl2 expression across neuronal cell types (R=0.62; Fig. 7F), and developmentally regulated (Fig. 7G). The cell type and developmental stage-specific splicing is due, at least in part, to splicing activation by Mbnl2 by binding to the downstream intron, as depletion of Mbnl2 and its homolog Mbnl1 dramatically reduces exon inclusion towards the level observed in GABAergic neurons (PSI=0.83 to 0.56; Fig. 7H). Interestingly, inclusion or skipping of this exon generates protein isoforms with distinct roles in neurite outgrowth (44), and therefore, may contribute to morphological and functional differences between glutamatergic and GABAergic neurons. These data together support the notion that Mbnl2 plays a pivotal role in driving the glutamatergic neuron-specific splicing program during brain development.

**Figure 7:**
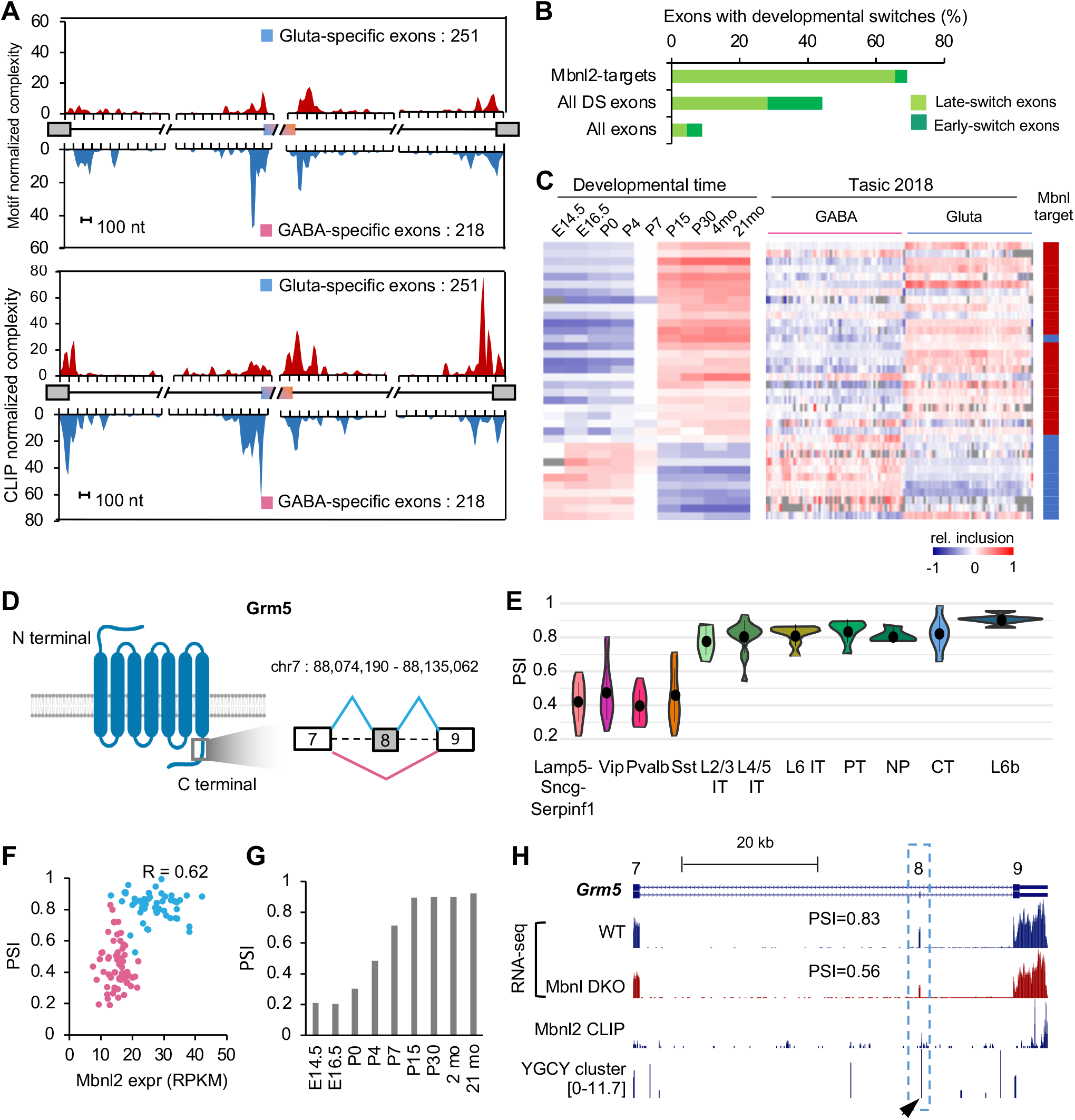
Mbnl2 drives a glutamatergic neuron-specific splicing profile. **A**, RNA maps showing enrichment of Mbnl binding sites (predicted YGCY clusters at the top and Mbnl2 CLIP tags at the bottom) in the upstream intron of cassette exons with GABAergic neuron-specific inclusion and in downstream intron of cassette exons with glutamatergic neuron-specific inclusion. **B**, DS exons regulated by Mbnl are enriched in late-switch exons during cortical development as compared to all DS exons or all cassette exons. Exons showing early or late developmental splicing switches were obtained from ref.(18). **C**, Comparison of splicing profiles of DS exons regulated by Mbnl during cortical development (18) (left panel) and across 117 neuronal cell types from Tasic 2018 (right panel). Exons activated or repressed by Mbnl are indicated in red and blue, respectively, in the scale bar on the right. **D-H.** Regulation of *Grm5* exon 8 by Mbnl2 is correlated with glutamatergic neuron-specific inclusion. **D**, Schematic diagram of Grm5 protein domain architecture and the region encoded by exon 8 with preferential inclusion in glutamatergic neurons. **E**, Violin plots showing Grm5 exon 8 inclusion in glutamatergic and GABAergic neuronal subclasses. Median and the interquantile range are indicated. **F**, Correlation of exon inclusion and Mbnl2 expression across 117 neuronal cell types. Glutamatergic and GABAergic neuronal cell types are colored in blue and pink, respectively. Pearson’s correlation coefficient is indicated. **G**, Grm5 exon 8 inclusion during cortical development. **H**, Mbnl2 bind to the downstream intron of Grm5 exon 8, as shown by Mbnl2 CLIP data and predicted Mbnl-binding YGCY clusters (arrowhead). Exon inclusion in the cortex is reduced upon depletion of Mbnl1/2 (Mbnl DKO).

### Graded, highly combinatorial splicing regulation across diverse neuronal cell types

We noted that ~46% (810 of 1,748) of neuronal class or subclass-specific DS exons show evidence of differential splicing at multiple hierarchical levels (Figs. S8A and Table S4; see examples in Fig. 4D). In parallel, over half (49/86) of differentially expressed RBPs showed differences between neuronal subclasses defined at multiple hierarchical levels (Figs. S8B and S9). We argue that the rich dynamics of RBP expression and downstream AS events across neuronal cell types can be used to infer the activity of an RBP in driving differential splicing in specific neuronal subclasses. Target exons activated by an RBP will show positive correlation with RBP expression, while target exons repressed by the RBP will show negative correlation. Therefore, the impact of the RBP in regulating global differential splicing across all or subsets of neuronal cell types can be evaluated using a gene set enrichment analysis (GSEA) (45) that examines the nonrandom distribution of RBP targets among all exons ranked by their correlation with RBP expression.

We applied GSEA analysis to 16 RBPs (from six RBP families) whose splicing targets were defined as described above. Strikingly, GSEA analysis revealed significant contribution of 9 RBPs, such as Mbnl2, Qk, and Nova1, and many of them across neuronal subclasses at multiple hierarchical levels (Fig. 8A,B). For example, Mbnl target exons show non-random distribution when ranked based on their correlation with Mbnl2 expression across all neuronal cell types, with activated exons at the top (positive correlation) and repressed exons at the bottom (negative correlation) (Fig. 8B). This is consistent with our analysis above indicating Mbnl2 as a major regulator of differential splicing between glutamatergic and GABAergic neurons. Importantly, similar patterns were observed when GSEA analysis was applied to GABAergic and glutamatergic neuronal cell types separately, suggesting that Mbnl2 also has a global impact on driving splicing dynamics within the two major neuronal classes. Indeed, we found 51 Mbnl target exons with differential splicing between L2/3 IT neurons and L6b neurons, which have the highest and lowest Mbnl2 expression within glutamatergic subclasses, respectively (Fig. 8C). Among them is GABA receptor gamma 2 subunit (*Gabrg2*) exon 9, whose inclusion is activated Mbnl2 by directly interacting with a downstream intronic binding site (18). Inclusion of this exon is positively correlated with Mbnl2 expression (R=0.76), with the lowest inclusion observed in L6b as compared to the other glutamatergic subclasses (Fig. 8D). In contrast, *Spint 2* exon 4, which is repressed by Mbnl most likely through exonic and upstream intronic binding sites, shows negative correlation with Mbnl2 expression (R=-0.72), with the highest inclusion in L6b neurons (Fig. 8E).

**Figure 8:**
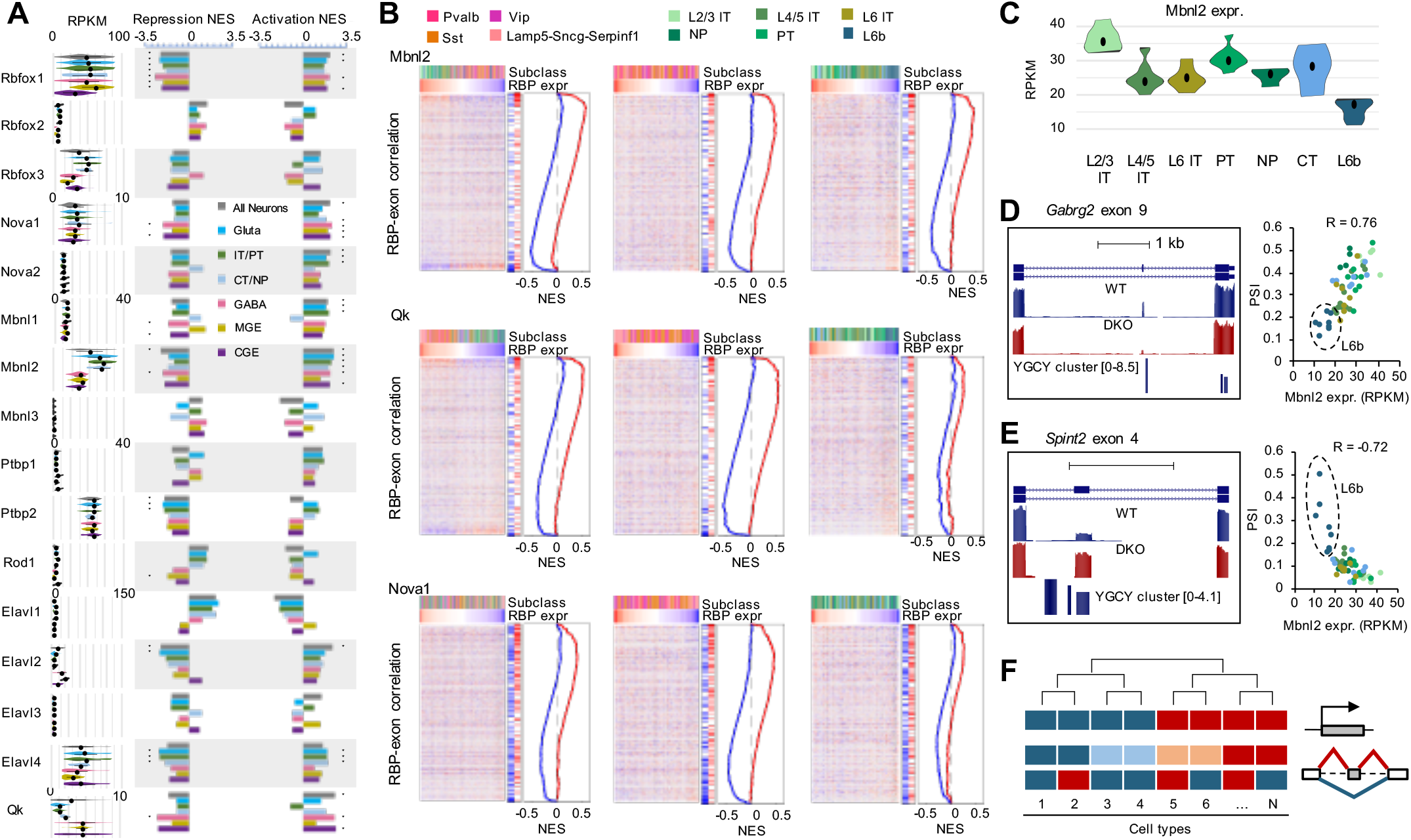
GSEA analysis of graded splicing regulation across diverse neuronal subtypes. **A**, RBPs with significant contribution to splicing regulation across neuronal cell types in various subclasses in are shown by the normalized enrichment score (NES) by gene set enrichment analysis (GSEA). Exons activated and repressed by each RBP family are analyzed separately. * FDR≤0.01 and NES≥1.8. **B**, GSEA plots of representative RBPs (Mbnl2, Qk and Nova1) across all neuronal cell types (left), across GABAergic cell types (middle) and across glutamatergic cell types (right). In each plot, exon inclusion across cell types is shown in the heatmap, in which exons are ranked by their correlation with RBP expression and the neuron types are ordered by the corresponding RBP expression in decreasing order. The color scales of correlation and RBP expression are shown on the left and top of the heatmaps. The neuron subclasses are indicated at the top using different colors. The repressed and activated targets of the corresponding RBP is indicated on the right of the heatmap together with the NES plots. **C**, Differential expression of Mbnl2 between L2/3 IT and L6 IT neurons from Tasic 2018 dataset. **D**, *Gabrg2* exon 9 activation by Mbnl2 through downstream intronic binding sites results in differential splicing across glutamatergic neuronal types. Tracks show exon inclusion in WT and Mbnl1/2 DKO cortices and predicted Mbnl-binding YGCY clusters (left). The correlation of exon inclusion and Mbnl2 expression across glutamatergic neuronal types is shown on the right. **E**, *Spint2* exon 4 repression by Mbnl2 through exonic and upstream intronic binding sites results in differential splicing across glutamatergic neuronal types. See legends of (D) for more detail. **F**, A schematic of the proposed model showing graded AS regulation that does not always conform to the hierarchical taxonomy of neuronal cell types defined based on transcriptional profiles.

Intriguingly, in some cases GSEA analysis revealed the global impact of RBPs on differential splicing across neuronal cell types, but the pattern of dynamics cannot be readily explained by the hierarchical taxonomy. For example, Nova1 and Nova2 do not show clear differential expression between adult glutamatergic and GABAergic neurons or between subclasses in our analysis (except downregulation of Nova1 in Lamp5-Sncg-Serpinf1), and yet GSEA analysis detected a strong impact of Nova1/2 on the differential splicing dynamics across all neuronal cell types, or within glutamatergic (Nova2) or GABAergic neurons (Nova1) (Fig. 8A,B). Similar observations were made for Rbfox1 and Ptbp2 (Fig. 8A). These results suggest a complex picture of graded, highly combinatorial, regulation of splicing across diverse neuronal subtypes (Fig. 8F).

## Discussion

Here we report systematic investigation of neuronal cell type-specific AS and the underlying regulatory mechanisms in the adult mammalian brain using deep scRNA-seq data (6, 7). Compared to recent work using bulk RNA-seq of purified neuronal populations or ribosome-engaged transcripts, this study offers two unique advantages. First, the unprecedented sequencing depth affords accurate quantification of AS and identification of DS exons with high sensitivity. We identified a larger number of DS exons from our analysis of scRNA-seq with stringent criteria than bulk RNA-seq data (34–36). Additional comparison with DS exons recently identified from bulk RiboTRAP-seq by another group suggests that the vast majority of DS exons we identified were not reported previously (25), and yet the newly identified DS exons in general show reproducible splicing patterns between the two scRNA-seq datasets and other independent bulk RNA-seq data, suggesting reliability of our analysis (Fig. S10).

Most importantly, the in-depth sampling of a large number of single cells without cell types defined a priori allows analysis of over 100 neuronal cell types that were defined from global transcription profiles rather than single reporter genes, including those missed in previous studies (e.g., the Lamp5-Sncg-Serpinf1 subclass of GABAergic interneurons) or treated as a homogeneous group (e.g., different subclasses of glutamatergic neurons). The hierarchical analysis of neuronal classes or subclasses enabled by scRNA-seq data provided several insights into the complexity and regulation of the transcriptome at the splicing level. One characteristic feature of AS regulation that became clear from our analysis is that it is “graded” rather than dichotomic or on-off regulation more commonly seen at the transcription level. This is supported by differential splicing of many exons detected at multiple hierarchical levels (Fig. 3D and Fig. S8A). In particular, a group of exons show coordinated differential splicing between GABAergic and glutamatergic neurons, between NP and other subclasses of glutamatergic neurons, and between L6b neuronal subclasses with local and long-range projection (Fig. 4). This putative splicing program should be further assessed for its possible role in regulating neuronal morphology and types of synapses related to the range of neuronal projection for multiple neuronal subclasses.

Very little is currently known about neuronal cell type-specific AS controlled by RBP splicing factors. It was recently shown that Nova2 is enriched in excitatory neurons and has distinct binding profiles in excitatory vs. inhibitory neurons in the developing cortex and cerebellum. Neuronal class-specific depletion of Nova2 affected splicing of a large number of exons, including a subset with differential splicing between excitatory and inhibitory neurons (46). Another study demonstrated that Rbfox1 is preferentially expressed in MGE-over CGE-derived cortical interneurons (47). Conditional depletion of Rbfox1 in two subclasses of MGE-derived, PV and SST interneurons resulted in distinct defects in synaptic connectivity and network excitability in mice. Hundreds of Rbfox1-dependent exons were reported in PV and SST neurons with minimal overlaps, lending the authors to argue a unique role of Rbfox1 in each interneuron subclass. Interestingly, the differential expression of these two RBPs between glutamatergic vs. GABAergic neurons is diminished in adult cortex from our analysis.

By integrating unbiased *de novo* motif discovery, differential RBP expression, RBP target networks, position-dependent RNA-maps, and GSEA, our analysis highlighted several RBPs with global impact on driving the distinct splicing profiles of glutamatergic and GABAergic neurons. Mbnl2, Celf1/2, and Khdrbs3 have preferential expression and higher activity in glutamatergic than GABAergic neurons, and therefore splicing activation or repression by these RBPs results in a glutamatergic neuron-specific splicing profile. On the other hand, Elavl2 and Qk are more highly expressed and potent in GABAergic neurons, driving towards a GABAergic neuron-specific splicing profile. Together, these RBPs regulate about one-third of DS exons. Consistent with lack of differential expression, a limited, if any, impact of Rbfox and Nova RBP families was detected on the distinct splicing programs between adult glutamatergic and GABAergic neurons. The role of Khdrbs RBP family in regulating excitatory neuron-specific splicing was previously established through extensive studies of the alternative exon SS4 of neurexin genes (23, 32, 33). We found 10 of 13 Khdrbs3-dependent exons are differentially spliced between cortical glutamatergic and GABAergic neurons, suggesting a striking degree of cell type specificity of this splicing program. On the other hand, the involvement of the other RBPs in regulating neuronal cell type-specific AS has not been reported and is unexpected to some extent. For example, Mbnl2 regulates developmental splicing switches in the brain, which are disrupted in myotonic dystrophy, but the exact cell types in which it acts were unclear (18, 43). Qk was mostly studied in oligodendrocytes with an important role in regulating myelination(48) and more recently in embryonic neural stem cells affecting cellular differentiation (49). C elegans homlogs of Celf (UNC-75) and Elavl (EXC-7) were previously shown to regulate cholinergic and GABAergic specific splicing in worms (50), although their function has likely diverged in vertebrates due to the dramatic expansion of these RBP families. Our results suggest potential productive avenues to revealing the functional significance of neuronal cell type-specific splicing by careful examination of neuronal phenotypes caused by depletion of these RBPs using *in vitro* and *vivo* model systems.

Consistent with the proposed graded AS regulation, differential expression of RBPs are mostly quantitative, and frequently detected between neuronal classes or subclasses at multiple hierarchical levels (Figs. S8B and S9), lending power to GSEA analysis to infer RBP activity by correlating RBP expression and exon inclusion across neuronal cell types without explicitly imposing the hierarchical structure. By applying GSEA analysis to different subsets of neuronal cell types, we found that most RBPs have a global impact on the splicing dynamics at multiple hierarchical levels. The most surprising finding resulting from our GSEA analysis is probably the inferred activity of RBPs, such as Rbfox1, Nova1/2 and Ptbp2, even when they do not show clear differential expression pattern between neuronal subclasses. In particular, Rbfox1 explains the splicing dynamics across neuronal cell types better than Rbfox3, although the latter shows distinct expression between glutamatergic and GABAergic neurons. Expression of these RBPs varies across diverse neuronal cell types composed of more homogeneous cell populations, and such variation clearly drives dynamics of AS across these cell types through splicing activation and repression, yet the dynamics do not always conform to the hierarchical taxonomy established using transcriptional profiles (Fig. 8A,B). Taken together, this study suggests a model in which transcriptional diversification by AS in the adult cortex is not simply downstream of transcriptional regulation, but an orthogonal mechanism to specify neuronal cell type identity and function through complex and graded regulation (Fig. 8F).

## Acknowledgements

We thank members of the Zhang laboratory for helpful discussion about the project. This study was supported by grants from the National Institutes of Health (NIH) (R01NS089676 and R01GM124486 to C.Z.). High-performance computation was supported by NIH grants S10OD012351 and S10OD021764.

## Author contributions

HF and CZ conceived the study; HF, DFM, SC, and MGM performed data analysis; VM and CZ supervised the work; HF and CZ wrote the paper with input from all authors.

## Materials and Methods

### Compilation of RNA-seq data

A summary of RNA-seq data analyzed in this study is provided in Table S1. Raw single-cell RNA-seq data of Tasic 2016 and Tasic 2018 in FASTQ format generated by Allen Brain Institute were downloaded from NCBI Short Read Archive (SRA) under accession numbers SRP061902 and SRP150473, respectively. Three bulk RNA-seq datasets of GABAergic and glutamatergic neurons, as well as RNA-seq datasets upon depletion of several RBPs including Qk, Celf1, Nova2, and Khdrbs3 were obtained from separate studies (Table S1). Mbnl1/2, Rbfox and Ptbp2-dependent exons detected by RNA-seq data were obtained from previous studies (18).

### Analysis of RNA-seq data and quantification of AS

All RNA-seq data were mapped by OLego (v1.1.2) to the reference genome (mm10) and a comprehensive database of exon junctions was provided for read mapping (51). Only reads unambiguously mapped to the genome or exon junctions (single hits) were used for downstream analysis.

To quantify AS, we used a comprehensive list of annotated AS events. Inclusion of alternative exons (percent spliced in or Ψ) was then quantified based on the number of exon junction reads using the Quantas pipeline (http://zhanglab.c2b2.columbia.edu/index.php/Quantas), as we described previously (16). To reduce uncertainty in estimating Ψ, we required exons to have junction read coverage ≥20. Gene expression was quantified using the same pipeline. For all quantifications, biological replicates were combined. For the Tasic scRNA-seq datasets, we used neuronal types defined in the original papers, and pooled core cells that were assigned to each cell type for AS and gene expression quantification.

### Dimensionality reduction, visualization and clustering of single cells using splicing profiles

We performed t-SNE analysis of single cells based on the splicing profile of all cassette exons as measured in the Tasic datasets. Starting with the exon inclusion (Ψ) matrix of known cassette exons across all core cells, we filtered exons by requiring exon inclusion level to be quantifiable (i.e., junction read coverage ≥20) in ≥20% of core cells and excluded cells with <1,000 quantifiable exons. In total, 19,853 cells with 3,174 exons in the Tasic 2018 dataset and 1,301cells with 2,972 exons in the Tasic 2016 dataset passed this filtering. We then imputed the exons with missing values in each cell using the k-nearest neighbor method (k = 50). Principal component analysis (PCA) was applied to the remaining exons and cells and the top PCs that explained ≥ 60% (for Tasic 2016 dataset) or 80% (for Tasic 2018 dataset) of the variance were used to generate 2-dimensional embeddings for data visualization using t-SNE (perplexity = 50; Fig. 1A and Fig. S2A,B). The splicing profiles of single cells were further clustered based on the top PCs using FindClusters, a shared nearest neighbor (SNN) based clustering algorithm, within Seurat (52) to identify clusters (using top 9 PCs with a resolution of 0.6 for the Tasic 2016 dataset, and top 30 PCs with a resolution of 0.8 for Tasic 2018 dataset). We then counted the fraction of cells in each cluster that were assigned to the originally defined neuronal cell types (Fig. 1B and Fig. S2C).

### Detection of differentially spliced exons

Differential splicing of cassette exons was performed using the Quantas pipeline, as described previously (16). Briefly, we used both exonic and junction reads that support each of the two splice isoforms and evaluated the statistical significance of splicing changes in the two compared conditions using a generalized linear model (GLM) (53). The false discovery rate (FDR) was estimated by the Benjamini-Hochberg procedure (54).

To identify exons differentially spliced between neuronal classes or subclasses using the Tasic 2016 and 2018 scRNA-seq datasets, we pooled read counts from single cells assigned to each neuronal cell types and treated these cell types within a group as biological replicates. This allowed us to reduce sampling noise at the single cell level while accounting for consistency of splicing across neuronal cell types within each group, resulting in more accurate identification of differentially spliced exons compared to other methods of pooling individual cells. For the Tasic 2016 dataset, we compared glutamatergic and GABAergic neuronal cell types and defined the exons with the following criteria as significant: read coverage≥20, Benjamini FDR≤0.05, exon quantifiable in ≥50% of cell types and |ΔΨ |≥0.2. We found filtering based on fraction of quantifiable neuronal cell types is effective in minimizing false positives, as it imposed reproducible differences across cell types of the compared groups. The Tasic 2018 dataset consists of cells from two different cortical regions. We performed three glutamatergic vs. GABAergic neuron comparisons by separating ALM and VISp cells and aggregating cells from the two regions. For this and following hierarchical analyses, CR Lhx5 and Meis2 Adamts19 types were excluded because they do not aggregate with the other glutamatergic and GABAergic neuronal types (6). We identified exons that are significant in any of the three comparisons using the same criteria as those used for the Tasic 2016 dataset. The final list of differentially spliced exons is the union of exons detected from the 2016 and 2018 datasets. We also reported the number of exons with less stringent cutoff |ΔΨ |≥0.1 while keeping all other criteria the same. For comparison, we performed differential splicing analysis between glutamatergic and GABAergic neurons using three bulk RNA-seq data. Exons with read coverage≥20, Benjamini FDR≤0.05 and |ΔΨ |≥0.2 was reported as significant.

To identify exons with neuronal subclass-specific splicing patterns at different hierarchical levels, we compared MGE vs. CGE originating GABAergic interneurons and detected differentially spliced exons using the same criteria as described above, except that we required | ΔΨ |≥0.1. We also identified exons differentially spliced between four major GABAergic subclasses Pvalb, Sst, Vip, and Lamp5-Sncg-Serpinf1, and between five major glutamatergic subclasses L6b, L6 CT, L5/6 NP, L5 PT, and IT. IT neurons were further subdivided into VISp L6 IT, ALM L5/6 IT, VISP L5 IT, ALM L2/3 IT and VISp L2/3 IT. For these analyses involving multiple neuronal subclasses, we performed one vs. other comparisons (e.g., Pvalb vs. the other non-Pvalb GABAergic neurons). If an exon is significant in more than one comparison (e.g., significantly higher inclusion in Pvalb vs. the other and also significantly lower inclusion in Vip vs. the other), it was assigned to the comparison with the smallest FDR.

### Annotation of differentially spliced exons

To assess the potential functional significance of glutamatergic vs GABAergic neuron-specific exons and accuracy of our computational pipeline, we annotated exons based on multiple types of characterization, including whether the alternative exon produces alternative protein products or cause nonsense-mediated decay (NMD), conservation of the AS pattern between mouse and human and/or rat, and whether the splicing pattern is neuron- or developmental stage-specific. These annotations were obtained from previous studies (16, 18) (Fig. S4A).

### Gene ontology (GO) analysis

GO analysis was performed using the 368 genes containing exons with differential splicing between glutamatergic and GABAergic neurons using the online tool DAVID (55). The background gene list for comparison is composed of all genes containing cassette exons with a sufficient read coverage in the cortex (coverage≥20). GO terms with Benjamini-Hochberg FDR≤0.05 were used to generate the network view using the “Enrichment map” application (56) in Cytoscape (v3.7.2) (57) (Fig. S4B).

To identify the shared and unique GO terms enriched in genes with differential splicing in different neuronal subclasses, we complied the gene sets with neuronal subclass-specific exons and followed with GO-Elite, a tool designed to identify a nonredundant set of GO terms and pathways for a particular set of genes (58). GO-Elite allows one to input multiple gene lists and summarize the overrepresented terms or pathways for each gene list. The significance of GO term enrichment was calculated by permutation (N=2000). GO terms with Benjamini-Hochberg FDR≤0.05 in at least one gene set were kept and the enrichment z-scores of these terms were used for visualization in the heatmaps (Fig. 3C).

### Identification of differentially expressed RBPs

A list of 393 previously annotated RNA-binding proteins (RBPs) was downloaded from RBPDB (59). We were able to extracted the RPKM expression values of 372 the RBPs and performed centroid linkage hierarchical clustering of these genes and neuronal cell types using median-centered log2(RPKM) values from the Tasic 2018 dataset. Differential expression analysis was performed using edgeR (60) incorporated in the Quantas pipeline. A gene is called differentially expressed between the two compared groups with fold change≥1.5, FDR ≤0.05 and the RPKM ranked in the top half of all the genes in at least one of the two compared groups. To identify RBPs with differential splicing between neuronal subclasses, we performed one vs. the other comparison (e.g., Vip vs. the other non-Vip GABAergic neurons). RBPs were assigned to the subclass with the lowest FDR if significant in multiple comparisons.

### *De novo* motif enrichment analysis

We performed k-mer (k=4, 5, 6) enrichment analysis using upstream or downstream flanking intronic sequences (200 nt in each region) and the exon sequence of glutamatergic neuron-specific and GABAergic neuron–specific exons; matched regions of all cassette exons with sufficient read coverage in the cortex (coverage≥20) were used as a control. For this analysis, repeat masked sequences were extracted, and the enrichment of each k-mer was evaluated using a hypergeometric test. The significantly enriched k-mers were identified with Benjamini-Hochberg FDR ≤ 0.05 in at least one of the six regions associated with glutamatergic neuron- or GABAergic neuron-specific exons.

Significant k-mers were then clustered based on both sequence similarity and their enrichment patterns in different regions associated with glutamatergic neuron- or GABAergic neuron-specific exons. For each pair of k-mers, sequence similarity was measured by a Pearson correlation coefficient calculated using Stamp (61); similarity of enrichment patterns was evaluated using the uncentered Pearson correlation of the six enrichment z-scores associated with glutamatergic neuron- or GABAergic neuron-specific exons. K-mer pairs with sequence similarity>2.67 (corresponding to 80% percentile of all k-mer pairs) and enrichment similarity >0.8 were connected to generate a k-mer graph for visualization by Cytoscape (v3.7.2) (57). Enrichment z-scores of representative k-mer clusters were visualized using heatmap.

### Analysis of target exons regulated by specific RBPs

Exons directly regulated by Nova, Rbfox, Mbnl and Ptbp were previously identified using an integrative modeling approach (18). In brief, this approach employs a Bayesian network to weigh and combine multiple types of data, including evidence of protein-RNA interactions as determined by CLIP data and bioinformatics predictions of motif sites, evidence of RBP-dependent splicing as determined by RNA-seq or microarrays, and several evolutionary signatures related to regulated AS. In addition, we also analyzed published RNA-seq data from Qk wild type and knockdown (KD) neural stem cells (49), Qk wild-type and knockout brains (62), Khdrbs3 wild type and knockout brains (33, 63) and Celf1 wild-type and knockout cardiac ventricles (64). Qk, Khdrbs3 and Celf1-dependent cassette exons was identified by requiring read coverage≥20, Benjamini FDR≤0.05 and |ΔΨ |≥0.1. Elavl-dependent exons were obtained from a previous study, which compared wild type and Elavl3/4 double knockout mouse cortices using exon-junction microarrays (|ΔI_Rank_|≥6.5) (65). These exons were intersected with glutamatergic neuron- and GABAergic neuron-specific exons, as shown in Fig. 6A.

To evaluate the impact of RBPs in regulating the global splicing profiles of neuronal cell types, we extracted the splicing profiles of the 469 DS exons between GABAergic and glutamatergic neurons in Tasic 2018 data. Missing values were replaced by KNN imputation using function impute.knn in R with k=10. PCA analysis were performed using function prcomp in R and neuronal cell types were projected to the two-dimensional space using the first two principal components (PCs) as shown in Fig. 6B. We then focused on RBPs with RNA-seq data derived from brain tissues and projected splicing profiles of wild type and RBP depletion samples to the same two-dimensional space.

### Inferring RBP activity in regulating neuron subclass-specific splicing using gene set enrichment analysis

Gene set enrichment analysis (GSEA) (45) was used to assess relationships between RBP expression and splicing patterns across all or subsets of neuronal cell types. Only exons with read coverage sufficient for splicing quantification in ≥50% of the cell types in each analysis were included. For a given RBP and set of neuronal cell types, exons were ranked based on the Pearson correlation between exon inclusion level and log2 RBP expression level. Cassette exons activated and repressed by several well-studied neuronal RBP families were obtained as described above, and served as the positive and negative “exon sets”. For each RBP, all cassette exons were ranked based on the Spearman rank correlation with splicing and log2 expression level of the RBP, and were used as input for the R package GSEAalong with the relevant “exon sets”. Exon set enrichment score was calculated using the option gsea.type=’preranked’. Normalized enrichment score (NES) and p-value was estimated based on 1000 random permutations of the input data, and the Benjamini-Hochberg procedure was applied for multiple test correction. We call significant enrichment by Benjamini FDR ≤0.01 and NES≥1.8.

## Notes

### Competing Interest Statement

The authors have declared no competing interest.

