## Supplemental Figures S1-10 for "Complexity and graded regulation of neuronal cell type-specific alternative splicing revealed by single-cell RNA sequencing"

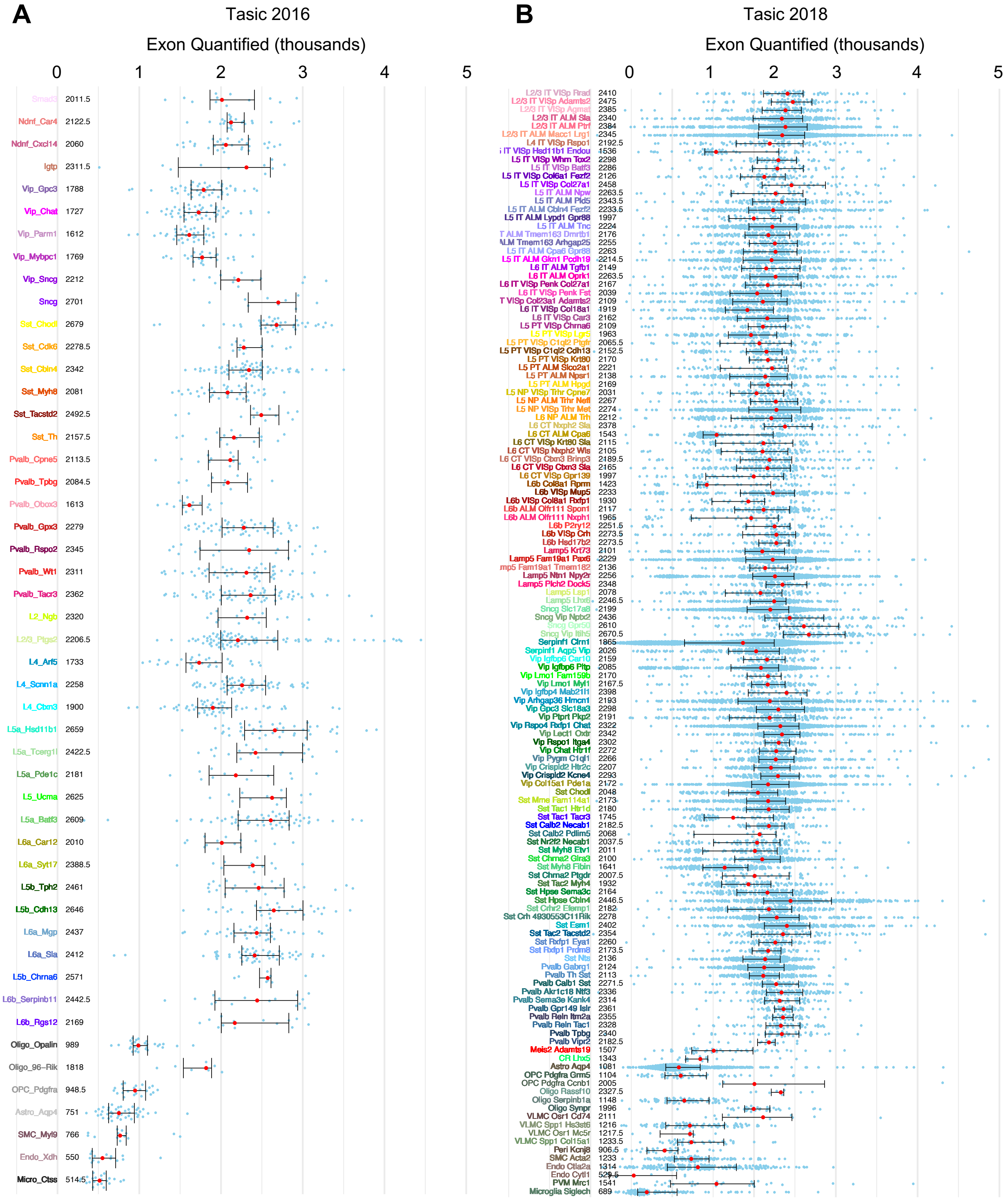

**Fig. S1: Splicing quantification for core cells segregated by transcriptional cell types. A**, Number of cassette exons with sufficient read coverage (≥20 junction reads) for quantification for all core cells in Tasic 2016 scRNA-seq data, grouped by cell types identified in the original paper. Red dot and whisker indicate the median and interquantile range. Median values in thousands of exons are also shown next to the cell type labels. **B**, Similar to (A), but for all core cells in Tasic 2018 scRNA-seq data.

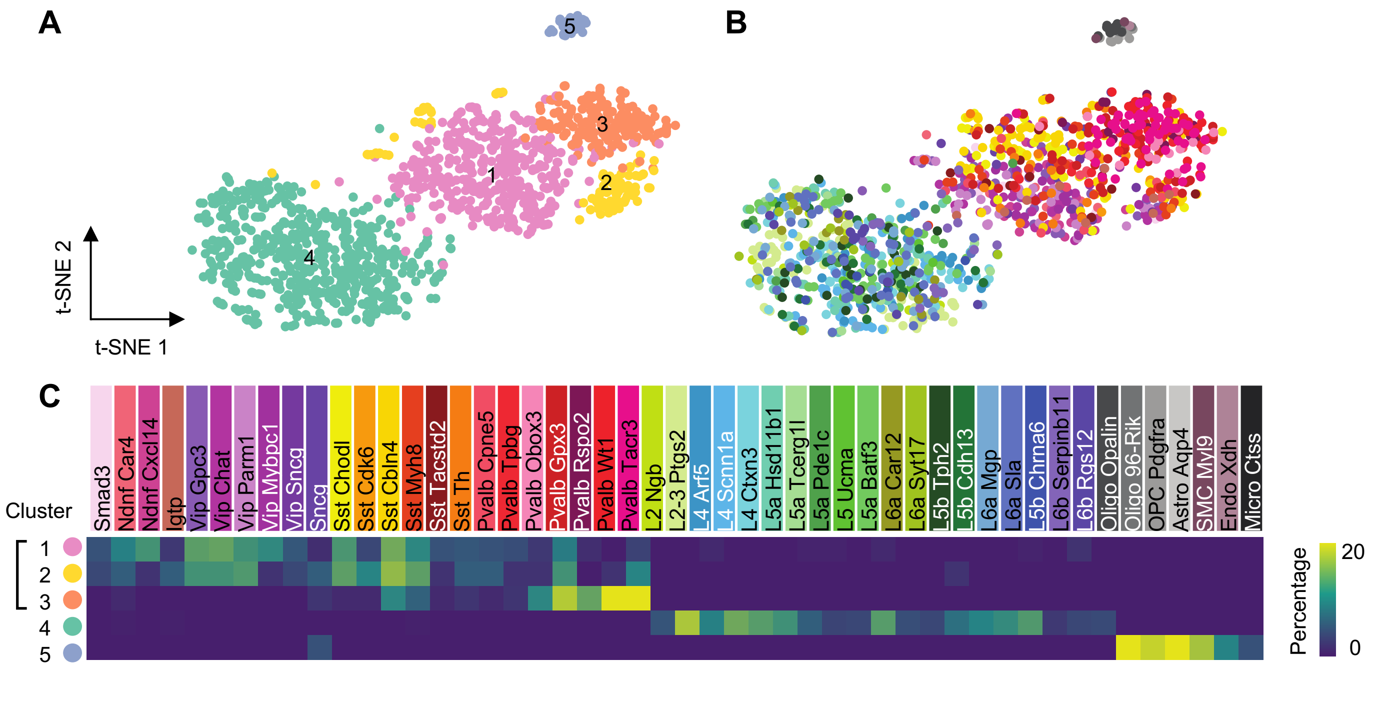


**Fig. S2: Splicing profile reveals major neuronal subclasses at single-cell level using Tasic 2016 dataset. A, B**, Two-dimensional t-distributed stochastic neighbor embedding (t-SNE) plots of 1,301 core cells. Cells are colored by clusters identified based on the splicing profile (A) or by the original cell types defined in the original study (B). **C**, Heatmap showing the overlap between the clusters defined based on splicing profiles and the original neuronal cell types defined based on expression profiles.

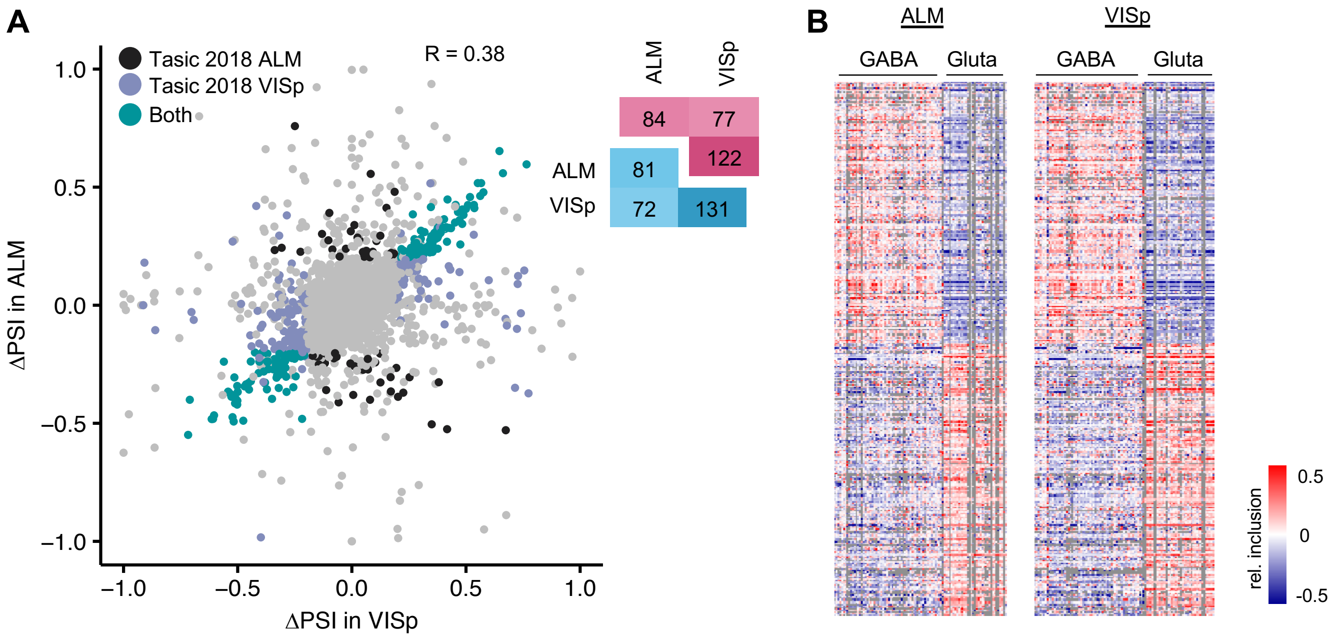


**Fig. S3: Splicing differences between glutamatergic neurons and GABAergic neurons detected in ALM and VISp are well correlated. A**, Comparison of ΔΨ (change in Percent Spliced In) between GABAergic and glutamatergic neurons of all cassette exons detected from ALM and VISp from the Tasic 2018 dataset. Black points represent the DS exons only detected from the ALM region, purple points represent the DS exons only detected from the VISp region, green points represent the overlap of DS exons detected in both regions, and grey points represent other non-DS exons. Pearson Correlation is indicated in the plot. The numbers of significant exons detected in each region are provided. Significant exons are defined by $|\Delta\Psi| \geq 0.2$ and FDR$\leq$0.05. **B**, Heatmap showing relative exon inclusion profiles (with median subtraction) of DS exons in glutamatergic and GABAergic neuronal cell types. The same list of DS exons as in Fig. 2C in the main text is shown.

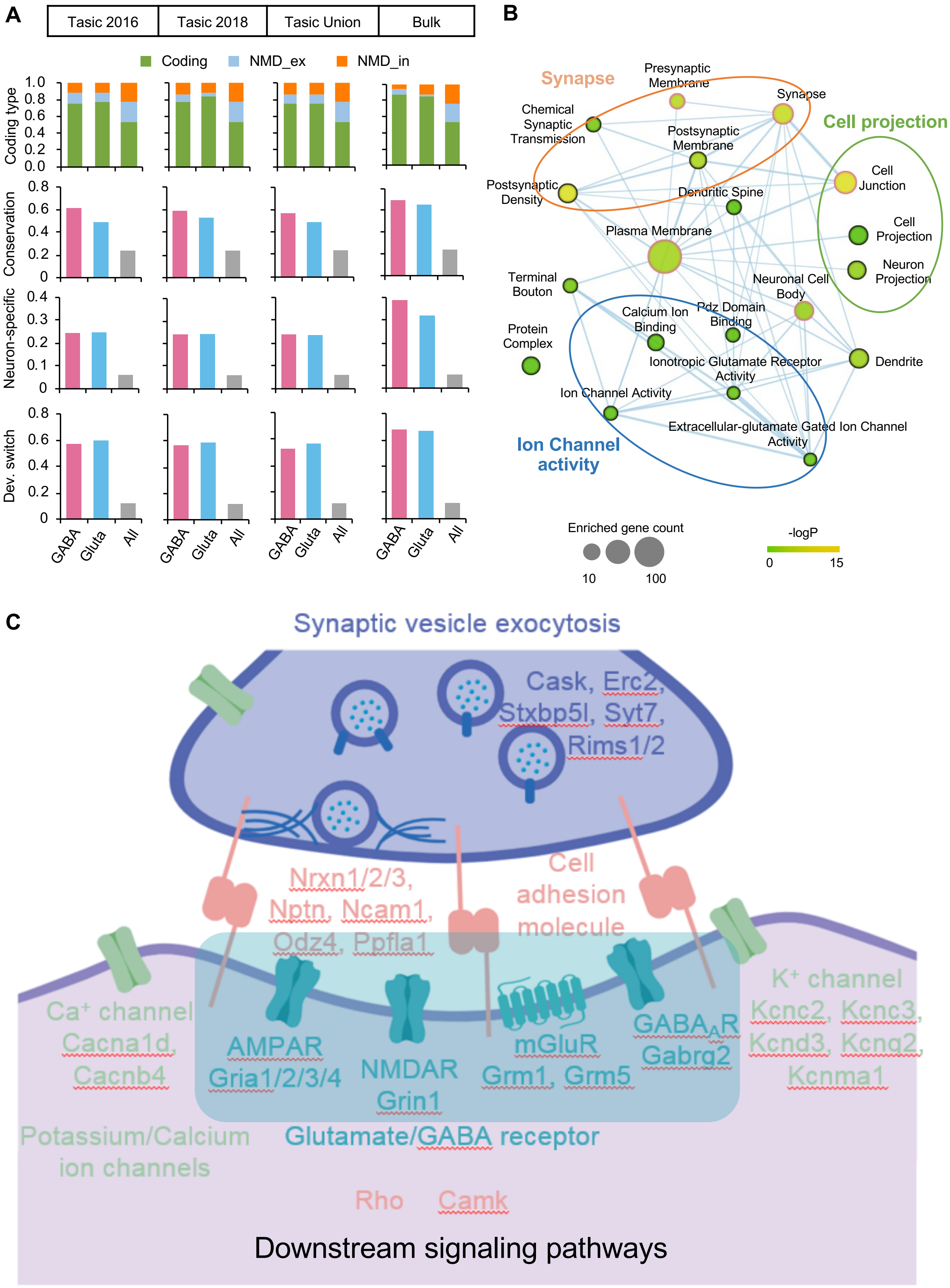


Fig. S4 (see legends on the next page)

**Fig. S4: Functional annotation of glutamatergic and GABAergic neuron-specific exons. A**, Annotation of DS exons detected in single cell and bulk RNA-seq data, as compared to all cassette exons used for control. The y-axis represents the proportion of DS exons in each category. **B**, Statistically enriched GO terms of genes with DS exons. The size and color represent the number and enrichment of genes associated with each term and related GO terms with overlapping genes are connected. **C**, Schematic illustration of the functional categories of genes with DS exons between glutamatergic and GABAergic neurons.

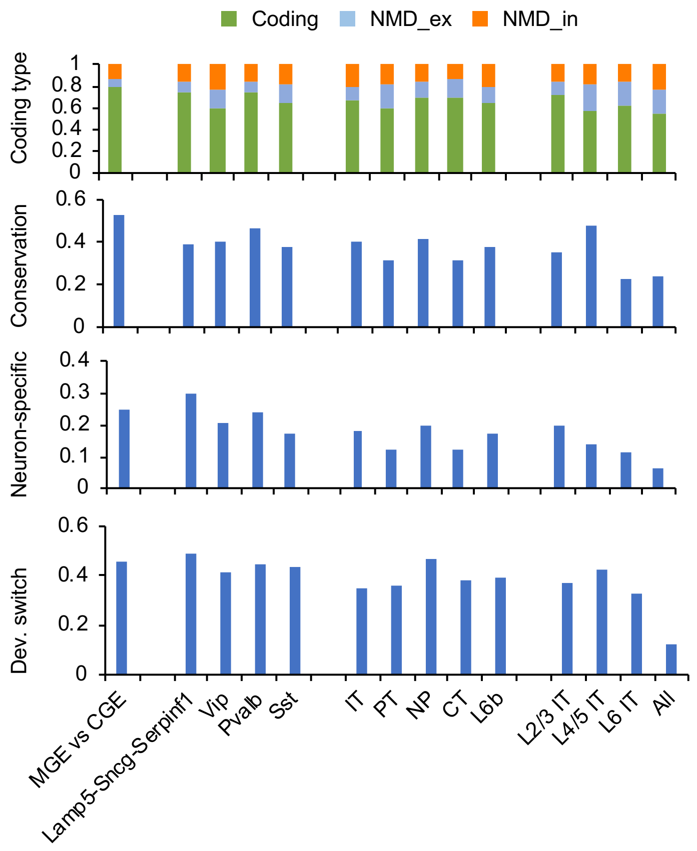


**Fig. S5: Characterization of neuron subclass-specific exons.** Annotation of DS exons detected at different hierarchical levels, as compared to all cassette exons used for control, is shown.

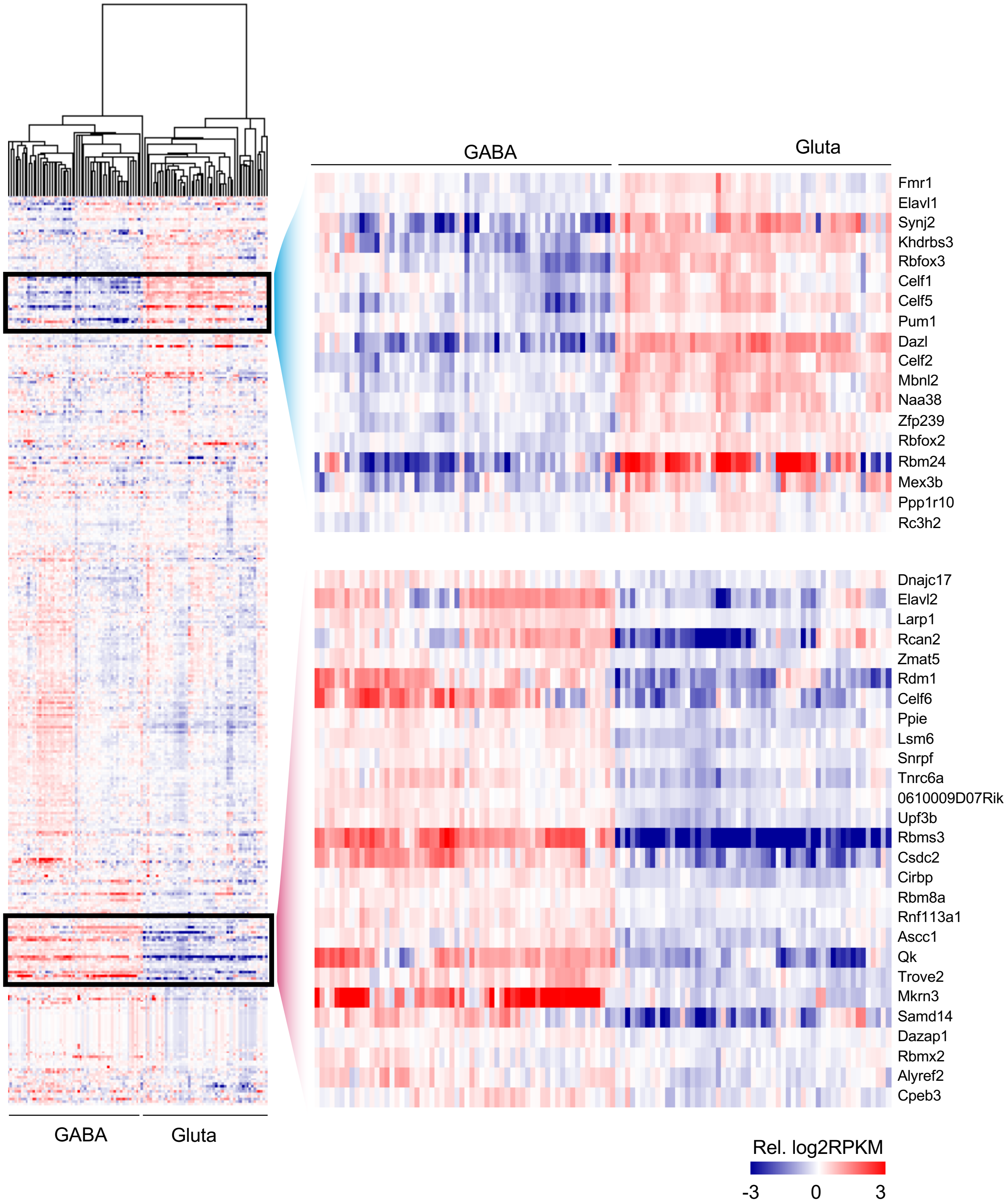


**Fig. S6: Unbiased clustering using RBPs expression profile separates glutamatergic and GABAergic neurons.** The heatmap shows the hierarchical clustering of gene expression profiles of 372 RBPs across the 115 neuronal cell types from the Tasic 2018 dataset. Note segregation of glutamatergic and GABAergic cell types. RBPs showing distinct expression pattern in the two neuronal classes are highlighted on the right.

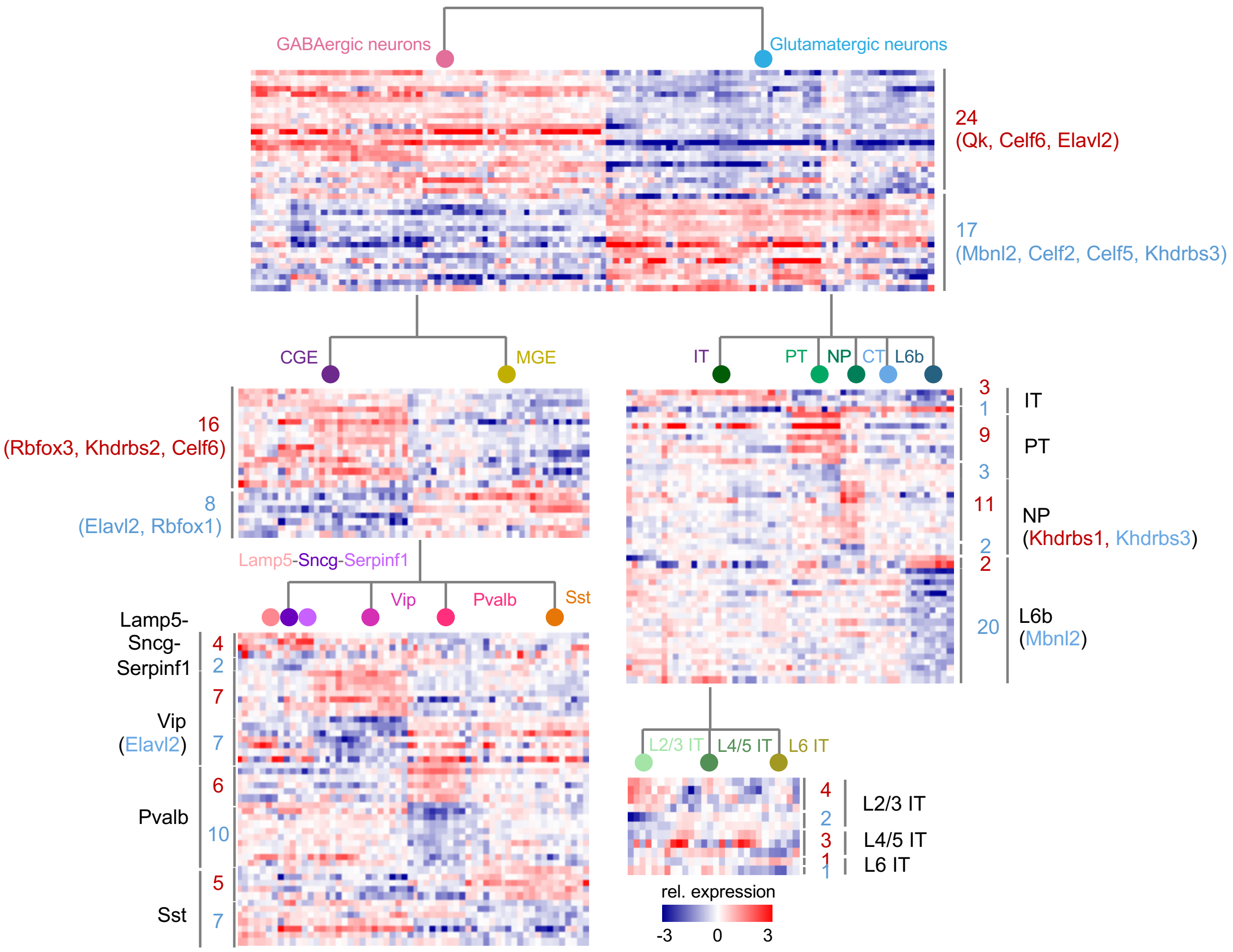


**Fig. S7: RBPs with differential expression between neuronal classes or subclasses at different hierarchical levels.** Heatmaps show expression pattern (log2 RPKM with median subtraction) of differentially expressed RBPs detected in different hierarchical levels. Each row represents an RBP and each column represents a neuronal cell type of indicated subclass. The numbers of differentially expressed RBPs as well as several representative RBPs detected in each comparison are labeled.

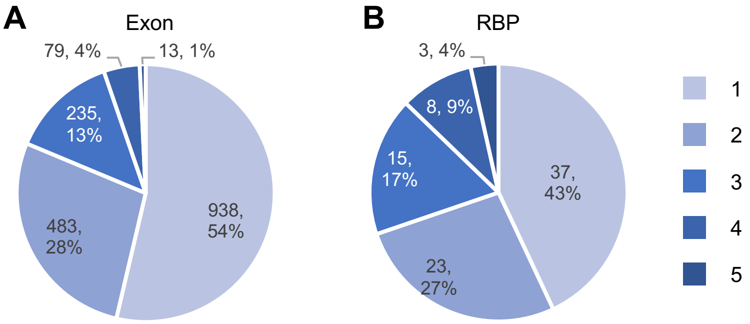


**Fig. S8: Graded, rather than dichotomic, differential splicing between neuronal subclasses. A**, Distribution of DS exons with respective to the number of comparisons contrasting different neuronal subclasses at different hierarchical levels, in which significant differential splicing is detected ($|\Delta\Psi| \geq 0.1$ and FDR$\leq$0.05 from Tasic 2018 dataset). **B**, Similar to (A), but for RBP with differential expression (fold change$\geq$1.5, FDR$\leq$0.05).

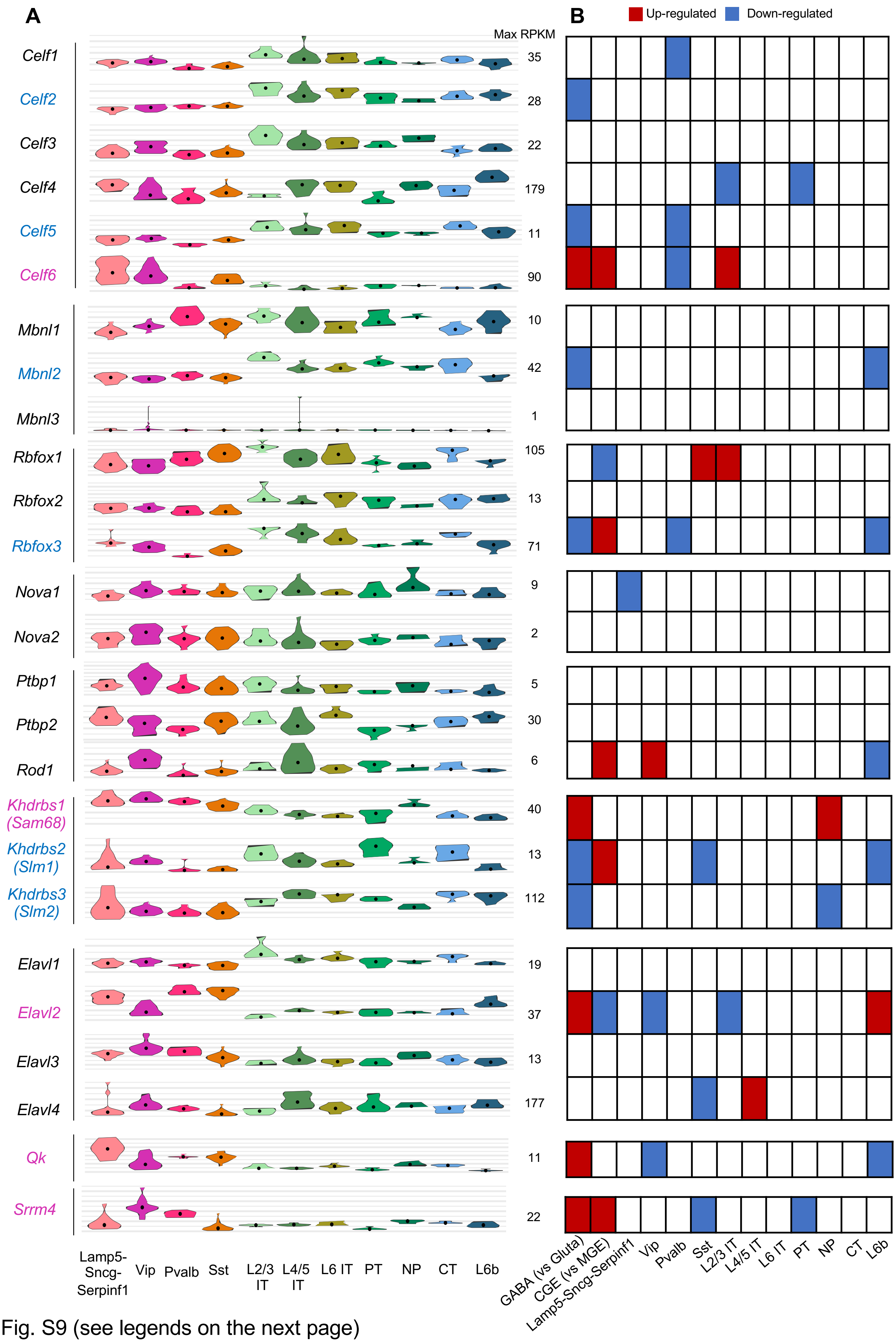


Fig. S9 (see legends on the next page)

**Fig. S9: Differential expression of RBPs previously known to regualte neuron-specific splicing.** The expression profiles of 9 RBP or RBP families across neuronal subclasses from Tasic 2018 dataset are shown on the left. Median and interquantile range of RPKM values of single cells in each subclass are indicated in each violin lot. Differential expression in 14 comparisons contrasting different neuronal subclasses is shown on the right. The RBPs with significantly higher or lower expression in the respective subclass are indicated in red or blue, respectively, in the table on the right.

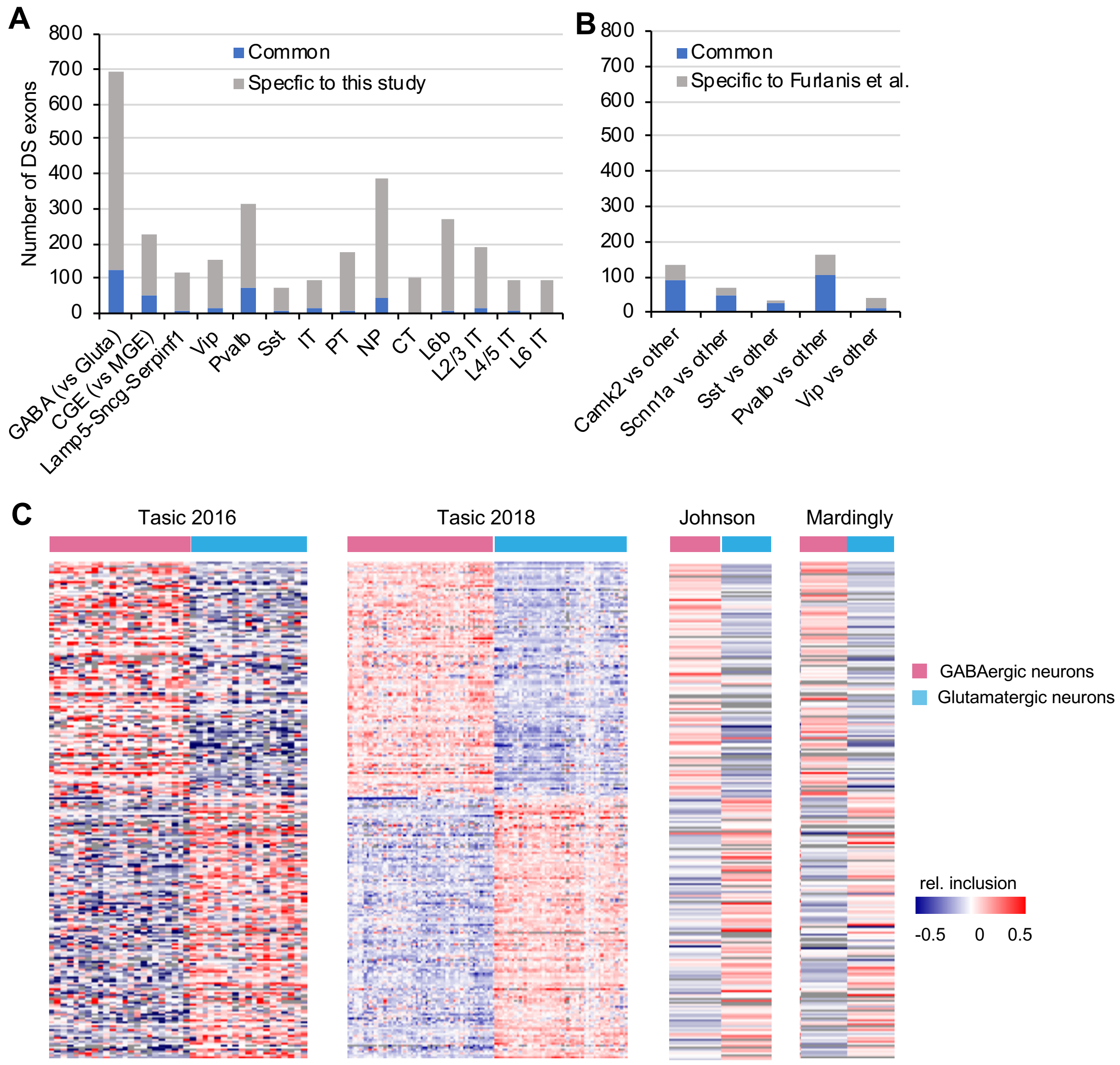


**Fig. S10: Comparison of DS exons detected in this study and those by Furlanis et al. A**, Number of common and unique DS cassette exons identified in this study compared with the lists from Furlanis et al. **B**, Number of common and unique DS cassette exons identified by Furlanis et al. compared with lists from this study. **C**, Heatmaps showing the splicing pattern of DS exons between glutamatergic and GABAergic neurons uniquely identified in this study.
